# Molecular Insights into the Rescue Mechanism of an HERG Activator Against Severe LQT2 Mutations

**DOI:** 10.1101/2024.03.13.584147

**Authors:** Amit Kumawat, Elisa Tavazzani, Giovanni Lentini, Alessandro Trancuccio, Deni Kukavica, Marco Denegri, Silvia G. Priori, Carlo Camilloni

**Affiliations:** Department of Biosciences, University of Milan, Milan, Italy; IRCCS Istituti Clinici Scientifici Maugeri, Pavia, Italy; Molecular Cardiology, Department of Molecular Medicine, University of Pavia, Pavia, Italy; Department of Pharmacy-Pharmaceutical Sciences, University of Bari Aldo Moro, Bari, Italy; Centro Nacional de Investigaciones Cardiovasculares Carlos III, Madrid, Spain

**Keywords:** hERG, LQT2, ICA-105574, molecular dynamics

## Abstract

**Background:** Mutations in the HERG potassium channel are a major cause of long QT syndrome type 2 (LQT2), which can lead to sudden cardiac death. The HERG channel plays a critical role in the repolarization of the myocardial action potential, and loss-of-function mutations prolong cardiac repolarization.

**Methods:** In this study, we investigated the efficacy and underlying molecular mechanism of ICA-105574, an HERG activator, in shortening the duration of cardiac repolarization in severe LQT2 variants. We characterized the efficacy of ICA-105574 *in vivo*, using an animal model to assess its ability to shorten the QT interval and *in vitro*, in cellular models mimicking severe HERG channel mutations (A561V, G628S, and L779P) to evaluate its impact in enhancing *I*_Kr_ current. Additionally, molecular dynamics simulations were used to investigate the molecular mechanism of ICA-105574 action.

**Results:** *In vivo*, ICA-105574 significantly shortened the QT interval. LQT2 mutations drastically reduced *I*_Kr_ amplitude and suppressed tail currents in cellular models. ICA-105574 restored *I*_Kr_ in A561V and G628S. Finally, *in silico* data showed that ICA-105574 stabilizes a pattern of interactions similar to gain-of-function SQT1 mutations and can reverse the G628S modifications, through an allosteric network linking the binding site to the selectivity filter and the S5P turret helix, thereby restoring its K^+^ ion permeability.

**Conclusions:** Our results support the development of HERG activators like ICA-105574 as promising pharmacological molecules against some severe LQT2 mutations and suggest that molecular dynamics simulations can be used to test the ability of molecules to modulate HERG function *in silico*, paving the way for the rational design of new HERG activators.

## Background

The *KCNH2* gene encodes a voltage-gated K^+^ channel (human ether-à-go-go, HERG) in neurons and cardiac cells. Its primary role involves cardiac repolarization through the rapid delayed rectifier K^+^ current (*I*_Kr_), which significantly influences the duration of the QT interval on the surface electrocardiogram (ECG) [1,2]. Abnormalities in the HERG channels are characterized by two distinct cardiac disorders, namely Long QT Syndrome (LQTS) and Short QT Syndrome (SQTS), which manifest as the prolongation or the shortening of the QT interval on the ECG. Congenital LQTS accounts for nearly 40% of cases [3,4], while the majority of LQTS cases stem from pharmacological blockade by various clinically used drugs. Mutations in HERG channels can result in a loss of function due to defective trafficking, abnormal channel gating or reduced ion permeation. Notably, approximately 90% of LQT syndrome type 2 (LQT2) mutants exhibit reduced synthesis and defective trafficking [5,6], yet certain mutations (e.g. G628S, N629D) [7,8] near the selectivity filter (SF) can maintain normal trafficking while failing to exhibit ion selectivity and conductance in physiological solutions. On the other hand, SQT type 1 (SQT1) mutations (e.g. N588K, T618I, S631A) are often associated with reduced or loss of inactivation, resulting in a shorter QT interval [9,10]. The large number of LQT2-associated missense mutations distributed throughout the HERG sequence poses a challenge to pharmacological treatments. Of note, Priori and coworkers showed that substrate-specific therapy with mexiletine, which shortens the QT interval, confers arrhythmic protection in LQT3 [11,12]. This suggests that, in principle, QT interval shortening is an effective strategy for the treatment of LQT2. This could be achieved with HERG activators [13], which enhance the rapid component of the delayed rectifier K^+^ current (*I*_Kr_) and may be able to shorten the duration of cardiac repolarization in LQT2.

Structurally, HERG channel is a homotetramer consisting of an N-terminal cytosolic Per-Arnt-Sim (PAS) domain followed by six transmembrane segments: S1-S4, forming the voltage-sensing domain (VSD), and S5-S6, together with the S5P turret region and the P helix, forming the pore-forming domain (PD); and the C-terminal cyclic nucleotide-binding homology domain (CNBHD), all intertwined by long disordered regions [14]. The HERG channel undergoes critical kinetic transitions between its closed (deactivation), active and inactivated states that regulate cardiac repolarization. Despite the recently available cryo-EM structure, the distinctive mechanism of rapid inactivation and recovery, and slow deactivation remains poorly understood due to the limited structural information available for the closed and inactivated states and are mostly the results of indirect indications obtained from homologues [15–20] . A recently determined cryo-EM structure for the HERG-astemizole complex has confirmed the general understanding that blockers bind to a hydrophobic pocket at the entrance of the SF, thus blocking the passage of K^+^ ions, and provides further tools for structure-based screening of HERG-related cardiotoxicity [21–24].

While HERG drug screening is primarily focused on the identification of drug-induced arrhythmic compounds, it has also revealed agonists (activators) that can enhance the outward current *I*_Kr_ amplitude [25]. These compounds show distinct mechanisms of action, with the majority inhibiting C-type inactivation, while others delay deactivation, increase channel open probability, or exhibit a combination these effects. From a structural perspective, these compounds appear to target different regions of the protein while avoiding blocking the channel [26]. Several activators have been investigated as potential candidates for different congenital LQTS mutations representing a novel approach to the prevention of ventricular arrhythmias associated with LQTS [26]. However, they carry the risk of inducing early repolarization, leading to an excessive correction of action potential duration (APD), which could ultimately shorten the QT interval to proarrhythmic levels (i.e., SQT). One such compound is ICA-105574, a derivative of 3-nitro-n-(4-phenoxyphenyl) benzamide, that is the most potent *I*_Kr_ activator to date [27,28]. Previous experimental and computational studies have revealed its binding site within a hydrophobic pocket located between two adjacent subunits, where it interacts with residues in the pore helix, the base of the SF, and the S6 segments [29–31]. ICA-105574 is proposed to act through a subtle change in the configuration of the selectivity filter that disrupts inactivation gating, with a demonstrated stoichiometric dependence where binding of multiple ICA-105574 molecules is required for optimal activity of the compound [32]. Importantly, ICA-105574 exhibits a complex mechanism of action, affecting both conductance and gating properties of the HERG channel [28].

In this study, we employed a multi-pronged approach using *in vivo* and *in vitro* electrophysiology and molecular dynamics (MD) simulations to explore the therapeutic potential and mechanism of action of ICA-105574 in LQT2 and in severe HERG mutations, specifically A561V, G628S, and L779P, derived from our cohort of LQT2 patients. First, we validated *in vivo* the ability of ICA-105574 to rescue QT prolongation in a sotalol-induced animal model mimicking LQT2. Then, we subcloned and expressed the above mentioned severe LQT2 HERG mutations in specific established stable cell lines to study their functional activity on current amplitude before and after ICA-105574 treatment, with our results indicating that ICA-105574 restores *I*_Kr_ amplitude in cellular models (A561V and G628S). Finally, we performed MD simulations to elucidate the molecular mechanisms of action of ICA-105574 in both wild-type and mutant HERG channels. There we revealed how ICA-105574 can rescue the permeability to K^+^ of the G628S mutation by stabilizing a pattern of interactions analogous to that observed in simulations of two SQT1 mutations (i.e., N588K and S631A) through an allosteric network connecting distant regions of the protein.

Our results allow us to propose that *in silico* methods are now sufficiently mature to evaluate molecules based on their ability to rescue channel conductance, indicate a molecular framework that can be used to rationalize its mechanism of action, and provide valuable insights into the therapeutic potential of ICA-105574.

## Methods

### ICA-105574 preparation

ICA-105574 was synthesized following a recently reported procedure by Zangerl-Plessl et al. [29], which is also employed in our recently published study [33], with only one modification - the use of HTBU (CAS number: 94790-37-1) in lieu of HATU (CAS number: 148893-10-1) as a coupling reagent.

### *In vivo* electrophysiological experiments

Male and female animals were obtained from Charles River Laboratories Italia SRL (Italy), and were kept in specific cages (Tecniplast, Italy), under controlled environment, with 12 hours light/dark cycle and room temperature at 23 °C. The guinea pigs had free access to water and food. After an acclimatizing period of one week, the animals were enrolled for the *in vivo* electrophysiological experiments. A subcutaneous telemeter was implanted to record the ECG signal in conscious and free-moving animals by the DSI PhysioTel Implantable Telemetry system. All procedures were performed in a quiet room to minimize animal stress. Each Guinea pigs was anaesthetized with 3-5% isofluorane-vet (Piramal Critical Care B.V. Netherlands). During the surgical operation, a thermally controlled heating pad was employed to avoid the drop of animal body temperature, and an ophthalmic tears solution was applied (Tears Naturale® P.M., Alcon) to avoid the ocular dehydration. A subcutaneous telemeter (PhysioTel ETA-F20 transmitter, Data Sciences International (DSI), USA) was implanted to record the ECG signal in conscious and free-moving animals by the DSI PhysioTel Implantable Telemetry system. After the surgical procedures, a recovery period was allowed to animals. In all animals, the baseline ECGs were recorded for 30 minutes in resting conditions in conscious animals. In analogy with previously published methodology [34,35] the LQT2 syndrome was chemically induced by administering 50 mg/Kg of sotalol per *os* into the cheek pouch [34], and the ECG signal was recorded for 90 minutes.

A single dose of ICA-105574 (15 mg/Kg, solved in: 50% Poly(ethylene glycol) 400, 30% N,N-Dimethylacetamide and 10% H_2_0) was intraperitoneally administered and the ECG signal was recorded for 90 minutes after injection. The ECG was acquired by Ponemha 6.50 software version (DSI), the measurement of ECG parameters was performed offline by 3 blinded investigators. To assess the trend of QTc change, while accounting for repeated measurements at pre-specified timepoints, we fitted a mixed effects linear regression model. Briefly, mixed effects linear regression models are a set of powerful statistical tools that permit the extension of linear regression to data with a hierarchical structure [36]. The QTc was thus inserted as the only outcome variable, and pre-set timepoints as the explanatory covariate. In all related analyses, p values were calculated using Kenward-Roger method of standard errors and degrees of freedom [37].

### Cloning of HERG-WT and mutants in heterologous systems

The commercial clone of full-length human HERG (NM_000238.3) cloned in pCMV6-XL4 (Origene) was subcloned in pmCherry-N1 (Clontech) to generate a functional chimera. Site-direct mutagenesis was performed using QuickChange II XL kit (Agilent Technologies) to generate the single HERG mutations: pmCherry-N1-HERG-A561V, pmCherry-N1-HERG-G628S and pmCherry-N1-HERG-L779P. All the plasmids were entirely sequenced. Primers are indicated in the supplementary information.

### Cell culture and transfection

To generate Human Embryonic Kidney (HEK293) stable cell lines, HEK293 cells were cultured in Dulbecco’s modified Eagle medium supplemented with 10% fetal bovine serum (FBS) and 1% penicillin/streptomycin, 1% of L-glutamine, sodium-pyruvate to 1mM and 1% non-essential amino acid solution at 37 °C in 5% CO_2_ and selected by G 418 Sulfate (Calbiochem). Cells were transiently transfected by Effectene (Qiagen) with 1µg plasmid (pmCherry-N1-hERG-WT and - A561V, -G628S and -L779P) and the HERG-mCherry-positive cells were sorted by BD FACS Aria III (Becton Dickinson)

### Immunofluorescence

HEK293 cells with pmCherry-N1-hERG-WT or mutants were stained with primary antibody anti-hERG (ab196301, Abcam), anti-Pan Cadherin (C1821, Sigma) or anti-Actin (PA116889, ABR). Secondary antibodies were DyeLight488-conjugated donkey anti-rabbit IgG (Jackson Lab). Dako mounting medium (Agilent Technologies) was applied to all slides. Confocal microscopy was performed with a Leica TCS-SP8 confocal microscope equipped with an HCX PL APO 40X/numerical aperture=1.25 oil immersion objective than exported to Adobe Photoshop (Adobe Systems, Mountain View, CA).

### Protein extraction and Immunoblotting

48 h after transient transfection of HEK293 cells with pmCherry-N1-hERG-WT or mutants (A561V, G628S or L779P) have been lysated in RIPA buffer (50 mM Tris HCl, pH 7.5, 150 mM NaCl, 1% NP-40, 0.5% sodium deoxycholate, 0.1% SDS) containing a cocktail of 10% protease inhibitors (Complete Mini, Roche). 30 μg of total proteins, quantified by the Pierce® BCA Protein Assay Kit (Thermo Scientific), were resolved by SDS-gel electrophoresis on Mini PROTEAN TGX Stain-Free 4-15% gradient Gels (Biorad) using Tris/Glycine/SDS buffer (Biorad), and blotted on 0.2 µm nitrocellulose using Trans Blot Turbo Transfer System (Biorad). The nitrocellulose membrane was saturated with a solution of TBS 1x, 0.1% Tween 20 (TBS-T) and 5% Skin Milk. After washes in TBS 1x + 0.1% Tween20 (TBS-T), the membranes were probed with different antibodies diluted in BSA 3% in TBS-T: anti-hERG (P0749 Sigma Aldrich, ab196301 Abcam) and anti-α Tubulin (2125 Cell Signalling) as reference protein. Secondary antibodies, conjugated with HRP (Promega), were diluted in BSA 3% in TBS-T and membrane was incubated for 1 hour at room temperature. Immunoblotting membranes were revealed using the Clarity Western ECL substrate (Biorad) and detected using ChemiDoc MP Imaging System (Biorad).

### *In vitro* electrophysiological experiments

Stable HEK293 cell lines expressing the single mutation were split and seeded to perform electrophysiological recordings. Only the cells expressing mCherry were chosen for current recordings by the Axiobserver microscope (Zeiss, Germany) equipped with the Colibri 7 (Zeiss, Germany) to recognize the fluorescent signal. The extracellular solution contained the following chemicals (in mM): 140 NaCl, 5 KCl, 1 MgCl_2_, 2 CaCl_2_, 10 HEPES, 10 Glucose and NaOH to reach a final pH of 7.4. A stock solution was prepared solving ICA-105574 by DMSO to reach a concentration of 10 mM. The stock solution was solved in extracellular solution to test ICA-105574 at the final concentration of 10 µM. The following intracellular solution (in mM): 130 KCl, 1 MgCl_2_, 10 HEPES, 10 EGTA, 5 Mg-ATP and KOH to stabilize the pH value at 7.2) was prepared [38]. The patch-clamp glass pipettes (WPI, USA) were pulled by a P-97 puller (Sutter Instruments, USA). hERG currents were recorded in voltage-clamp mode by the patch-clamp technique in whole-cell configuration. The Multiclamp 700B (Axon Instruments, USA) patch-clamp amplifier was connected to the DigiDadata 1322A (Axon Instruments, USA), an AD/DA converter related to a personal computer running pClamp software (Version 9.2, Axon Instruments, USA) used to record and analyse the hERG currents. Data were tested by two-way ANOVA with an appropriate post-hoc correction (Šidák test). Two-tailed P values were calculated with the statistical significance threshold set at p< 0.05.

### Molecular docking protocol for ICA-105574

ICA-105574 (3-nitro-N-[4-phenoxyphenyl]-benzamide) was docked into the cryo-EM structure of the hERG channel (PDB: 5VA2) in the WT using the induced fit docking (IFD) program from Schrodinger software (Release 2021-4, Schrödinger, LLC) [39]. The WT-ICA system was generated using this process, while the A561V-ICA and G628S-ICA systems were created by mutating specific residues in the WT-ICA system during the CHARMM-GUI system preparation protocol [40]. The binding site of the molecule was defined within the hydrophobic pocket situated below the selectivity filter (SF) between two adjacent subunits, as predicted by Zangerl-Plessl *et. al*. [29]. The binding site for each molecule was defined by a combination of residues from two neighboring subunits (chain-A: 622, 649; chain-B: 557, 652). The size of the box was set to 10 Å, with the geometric center being the centroid of the selected residues. Residues within a 5 Å radius of the binding site were set as flexible, except for the selectivity filter residues, which were constrained to prevent changes in the SF pore. Furthermore, it has also been suggested that multiple ICA-105574 molecules are necessary for channel activity [32]. Consequently, we repeated the docking protocol to place another ICA-105574 molecule into diagonally opposite fenestration. Once we generated and selected the docked structure with two molecules based on the docking score and glide gscore, we constructed the system with a POPC bilayer, water, and salt (KCl), followed by a brief equilibration phase with the protein restrained and the ligand flexible, ultimately generating the initial structure for the MD simulations.

### Molecular dynamics (MD) simulations details

#### (a) Tetrameric hERG channel

MD simulations were performed for eight different systems that represent WT (WT), two SQT1 mutations (S631A and N588K), two LQT mutations (A561V and G628S) and ICA-105574 bound WT (WT-ICA), A561V mutation (A561V-ICA) and G628S mutation (G628S-ICA). In the present study, the simulations were performed with the PAS domain unlike the earlier computational studies which include only the transmembrane region with the C-terminal domain. The open state cryo-EM structure of hERG was used for the WT (PDB: 5VA2) and SQT1 mutation S631A (PDB: 5VA3) [18]. hERG channel with SQT1 mutation N588K or LQT2 mutations A561V and G628S were generated by mutating the WT, while ICA-105574 bound systems were generated using the molecular docking protocol. The missing extracellular loops of the transmembrane domain (L432-L452, L510-L520, P577-I583 and W597-G603) in the cryo-EM structure were modelled using Modeller [41]. The membrane-protein systems were prepared using charmm-gui web service [40]. The protein was embedded in POPC bilayer and solvated with TIP3P water model [42]. On average, 400 POPC molecules were introduced into both the outer and inner side of the lipid bilayer. The solvated systems were neutralized by adding counterions and an appropriate number of K^+^ and Cl^-^ ions were added to maintain the salt concentration of 200 mM. A set of preliminary simulations with a mix salt concentration of 200 mM KCl and 200 mM NaCl were performed to ascertain K^+^ ion selectivity of channels in the present setup. The average box size of the resulting systems was ∼16.5 x ∼16.5 x ∼17 nm^3^. A constant electric field was applied to the system in the Z-direction to impose transmembrane voltage difference in the range +700 mV to +750 mV to observe outward potassium ion permeations. All simulations were performed using CHARMM36m forcefield [43] with NBFIX parameter [44] for potassium ion and carbonyl oxygen interaction. The parameters for ICA-105574 molecule were obtained using CGenFF program [45], which performs automated assignment of parameters and charges by analogy and compatible with CHARMM forcefield. Simulations (Tables S1 and S2) were performed with and without the application of a restraint on the backbone dihedral angles (φ and ψ) of the selectivity filter (SF) residues from S624 to G628 using alphabeta collective variable defined in PLUMED software [46]. The reference value for the dihedral angles were used from the cryo-EM structure. Potassium ions were placed in the SF at different sites (S1-S4) to prevent the collapse of the SF in the beginning of the simulations. Ion permeation mechanisms in potassium ion channels have been found to be ambiguous in nature, depending on the type of force field considered, the membrane potential, and the charge corrections for the ions. In preliminary simulations with dihedral restraints (Table S1B), we observed mostly a soft knock-on mechanism for the ion permeations using the CHARMM force field with NBFIX parameters, regardless of the initial configuration of the ions ([KKKKK] or [KWKWK]). In addition, the mixed KCl and NaCl simulations for the wild-type system established the ion selectivity with outward K+ ion movement and no inward/outward Na+ ion movement in the presence of torsional restraints. Therefore, subsequent production simulations were run with [KWKWK] ion configuration and KCl salt only for uninterrupted ion permeations (see Movie S1 and Table S2). All systems were simulated in two replicates with production run ranging from 1 to 2 μs for each system. The systems were energy minimized using steepest descent algorithm. This was followed by multiple step equilibration protocol in NVT and NPT ensembles with gradual decrease in the position restraints of protein and lipids (cf., Table S3). The temperature was maintained at 310 K using the velocity rescale method [47] with relaxation time of 1 ps and the pressure was kept constant at 1 bar using c-rescale barostat [48]. All simulations were performed using periodic boundary conditions and the long-range interactions were calculated using the particle mesh Ewald (PME) summation method [49]. The cut-off distance for electrostatic and van der Waals interactions was set to 1.2 nm. The bonds were constrained with LINCS algorithm [50], and the integration time step was set to 2 fs. All simulations were performed using GROMACS software (version 2021.X) [51] with PLUMED (Version 2.8) [46]. The stability of the simulated trajectories were ensured based on backbone RMSD computed over the production run for tetramer assembly and the pore domain of different subunits (Figure S1, S2).

#### (b) Cyclic-nucleotide binding homology domain (CNBHD) of hERG channel

MD simulations were performed for CNBHD to access the effect of LQT2 mutation L779P on the overall stability and conformation of the domain. The L779P mutation corresponds to the residue number 69 in the PDB file of the solution structure (PDB ID: 2N7G) [52]. All simulations were performed using GROMACS [51] with charmm36 forcefield [43] and the TIP3P water model [42]. The systems were solvated, neutralized by adding counter ions and an appropriate number of Na^+^ and Cl^-^ ions were added to maintain the salt concentration of 150 mM. This was followed by energy minimization using steepest descent method. The temperature was maintained at 310 K using v-rescale thermostat [47] and pressure was kept constant at 1 atm using c-rescale barostat [48]. The simulations were performed using periodic boundary conditions and particle mesh Ewald method [49] was used to treat long range electrostatic interactions. The cut-off distance for short-range electrostatic and van der Waals interactions was set to 1.0 nm. The bonds were constrained with LINCS [50]. Three independent simulations were run for 1 μs each with frames saved at every 10 ps.

## Results and discussion

### *In vivo* electrophysiological experiments demonstrate the shortening of the QT interval mediated by ICA-105574

To evaluate the efficacy of ICA-105574 in shortening the QT interval in LQT2 patients, we performed *in vivo* electrophysiological experiments by recording the ECG signal in conscious guinea pigs. Our choice was primarily due to different issues: (1) to enroll the animal model with the cardiac electrophysiological phenotype as closely as possible to human; (2) to avoid any possible effect of the anesthetic on the sensitivity to drug [53,54] ; (3) to observe any possible side effects in free moving animals; (4) to employ the smallest animal model with an easily recognizable T wave requiring the smallest amount of drug possible. To combine all the above reported aspects, we decided to employ the guinea pigs, since their cardiac ion currents are similar to humans [55] , with an easily identifiable P, Q, R, S and T waves recapitulating the LQT2 phenotype.

We investigated the effect of the *I*_Kr_ activator ICA-105574 in a chemically induced LQT2 guinea pig. The LQT2 condition was mimicked by pre-treating the guinea pig with 50 mg/Kg of sotalol, with a pronounced prolongation of the QTc interval from a baseline of 255 ms (IQR: 252-259 ms) was consistently obtained in all guinea pigs (ΔQTc +41 ms, +17%; <0.001) [34,56,57]. Despite marked QTc prolongation, during the electrocardiographic registration no spontaneous arrhythmic events were observed. Following the demonstration of successful induction of LQT2, a single intraperitoneal dose of ICA-105574 at 15 mg/Kg was administered. Immediately upon drug administration, QTc interval normalization was consistently obtained in all guinea pigs (ΔQTc -43 ms, -14%; <0.001). This effect was transient and by 30 minutes after administration QTc interval normalization waned with QTc lengthening back to values comparable to ones prior to ICA-105574 administration. No other biologically significant effects on the principal electrocardiographic parameters were observed (Figure 1).

**Figure 1.**
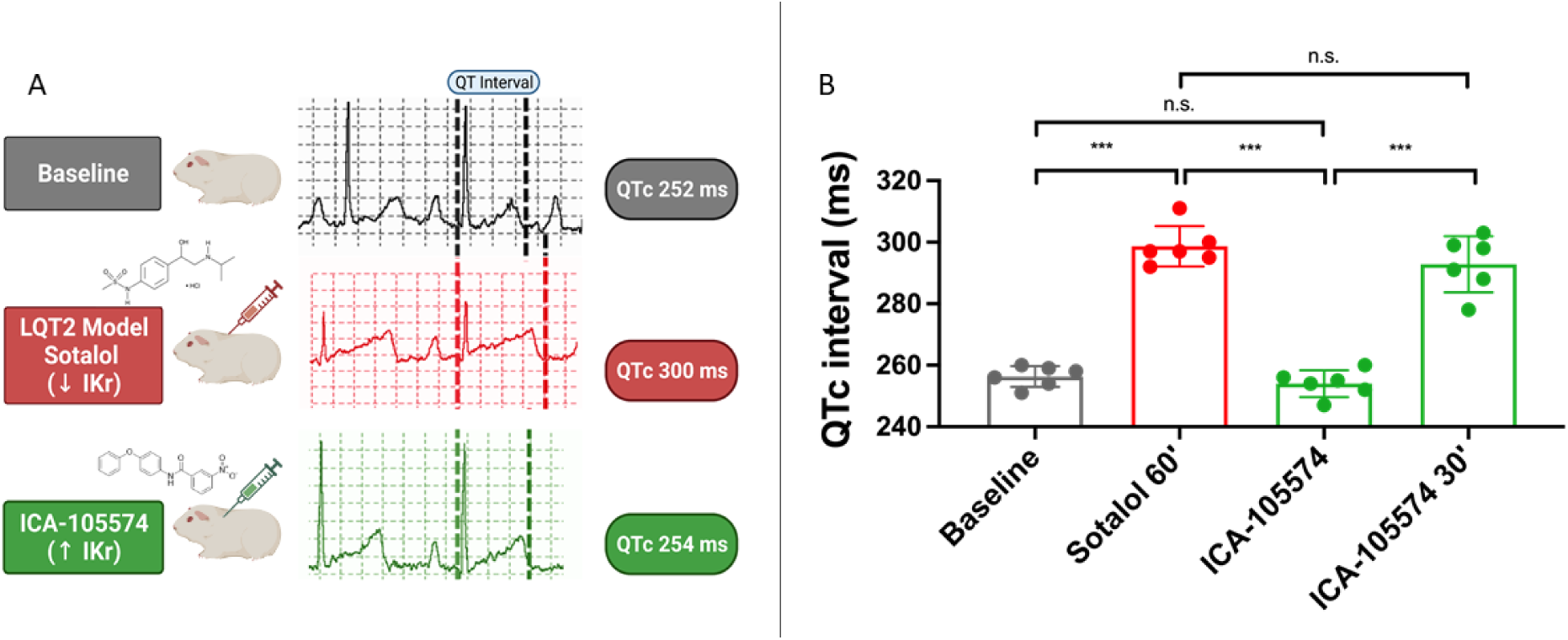
(A) *In vivo* LQT2 ECG recordings. LQTS was successfully induced in all animals after sotalol treatment (252±10 ms vs. 295±9 ms for the baseline and sotalol treated guinea pigs, respectively; p<0.001). Importantly, a pronounced and statistically significant shortening of the QTc interval was obtained in guinea pigs after ICA-105574 administration, normalizing the QTc in all animals (295±9 ms vs. 254±4 ms, before and after ICA administration, respectively). Importantly, despite the marked QTc shortening, no arrhythmic events were observed in the studied animals after ICA-105574 administration. (B) Pairwise comparisons were performed and the main relevant differences regarding the ECG values, are shown as histograms.

### *In vitro* electrophysiological analysis of ICA-105574 with, A561V, G628S and L779P

We then characterized the functional properties of HERG mutations (A561V, G628S and L779P) and their response to ICA-105574 as an *I*_Kr_ activator. We generated chimeric proteins (pmCherry-N1-HERG-WT, -A561V, -G628S and -L779P) and established single stable cell lines for each condition to perform electrophysiological studies following ICA-105574 perfusion. The pmCherry-N1-HERG-WT was transiently transfected into HEK293 cells and protein expression was assessed by Western blot and localized by confocal microscopy. These preliminary analyses confirmed the correct expression of the chimera and its distribution along the cell membrane (Figure S3). Similarly, we generated and performed the experiments for all the HERG mutations (Figure S4). Both mCherry-N1-HERG-A561V and -G628S were stained along the plasma membrane with some diffuse staining in the cytoplasm, similar to the HERG-WT. In contrast, the HERG-L779P was mainly localized in the cytoplasm, with enrichment around the perinuclear region. The localization of all HERGs proteins were not affected by ICA-105574 treatment.

To define the functional changes induced by ICA-105574 on LQT2 mutations, electrophysiological analysis was performed in HEK293 stable cell lines expressing HERG-WT and mutants (A561V, G628S, L779P). *I*_Kr_ was elicited by the voltage-clamp protocol showed in Figure S5 in control conditions and in presence of 10 µM ICA-105574. From a holding potential of -80 mV, voltage pulses were applied ranging from -60 to +70 mV in 10 mV increments, followed by repolarization to -60 mV. Using this activation protocol, it was observed that the HERG channels functionality was impaired, as the mutations led to suppression of the tail currents. To properly study the voltage dependence of activation, it is necessary to analyze the tail current amplitude elicited by a single repolarizing potential (–60 mV, Figure S5), which reflects the proportion of channels activated during the preceding depolarizing pulse. In the studied HERG mutations, suppression of the tail currents made it impossible to accurately calculate the activation curve parameters, including the half-activation voltage (V_1/2_) and the slope factor (κ), which are necessary to evaluate the voltage dependence of HERG channel activation induced by the mutations (as in Sanguinetti et al. for A561V and G628S [58]) and by ICA-105574. The A561V mutation reduced the steady state pre-pulse current amplitude and partially altered the HERG current behavior (Figure 2 and Figure S5), resulting in a slightly flattened activation bell-shaped curve, whereas G628S and L779P mutations caused a drastic reduction in the *I*_Kr_ current amplitude across all voltages (Figure 2 and Figure S5). The G628S currents decrease at negative voltages as the I/V curve approaches the equilibrium K^+^ value.

**Figure 2.**
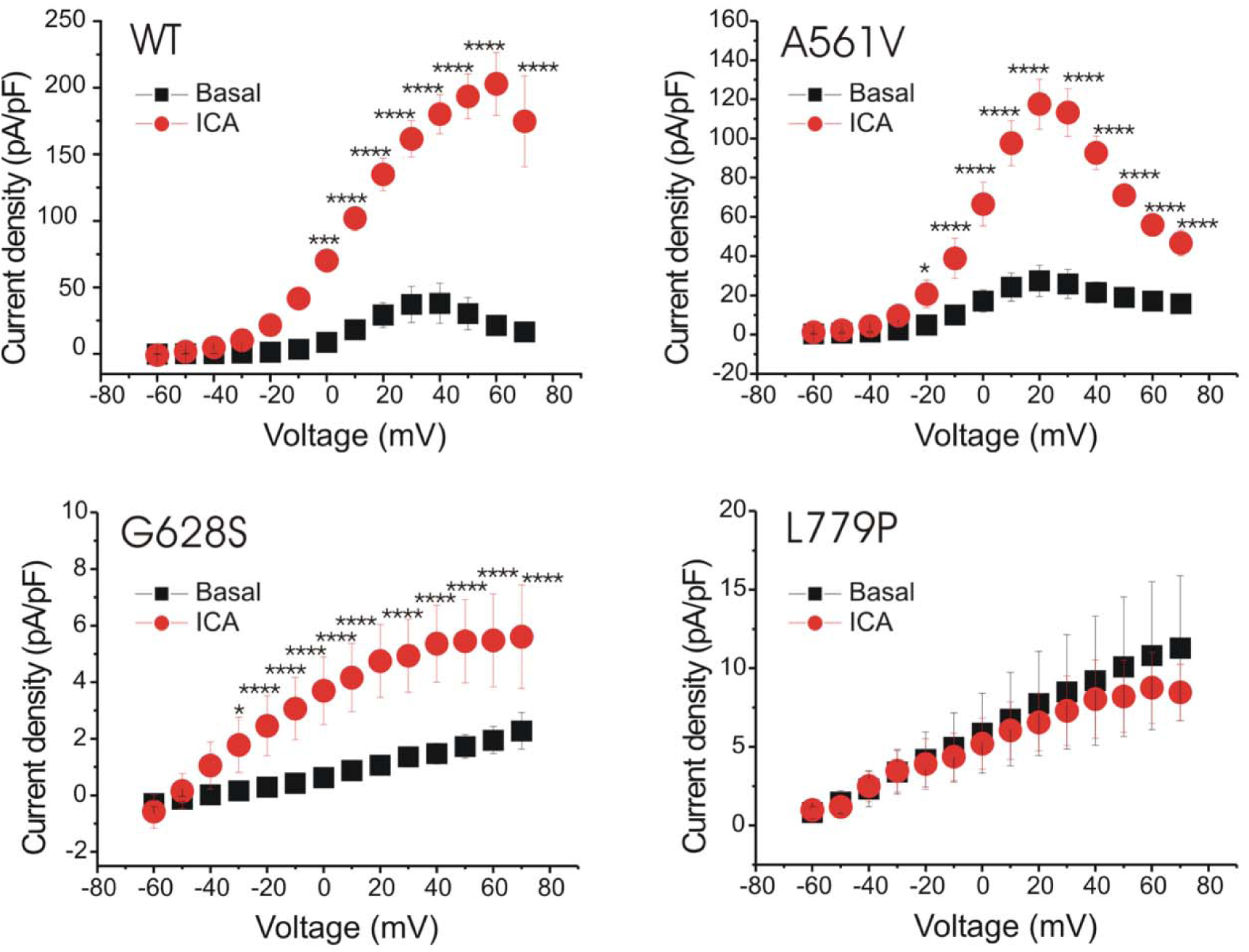
I/V relationships in HERG WT and mutants. The I/V graphs are derived by voltage clamp recordings showing WT, A561V, G628S and L779P activation currents recorded in basal condition (black square) and after 10 µM ICA-105574 perfusion (red dot). Each value is represented as mean□±□s.e.m. Statistical analyses were conducted using two-way ANOVA with Šidák post test. \**P*□<□0.05, \*\**P*□<□0.01, \*\*\**P*□<□0.001 and \*\*\*\**P*□<□0.0001. WT: n=5, A561V: n=6, G628S: n=6, L779P: n=4.

However, unlike the WT, the G628S channel remains open even at these negative voltages, indicating altered gating behavior. Additionally, the L779P mutations produced currents that failed to reach a peak or plateau phase, preventing the formation of a bell-shaped curve in the I/V activation relationship that is typical of WT. Taken together, these data allow us to speculate that G628S mutation, located in the pore region, negatively altered channel opening, consistent with previous findings for G628S [8,31,58,59], while L779P, located in a peripheral region, may exhibit a different mechanism of dysfunction, possibly related to protein misfolding, as also suggested by the observation that most of the protein is found in the cytoplasm, with enrichment around the perinuclear region (Figure S4). The HERG stable cell lines (WT, A561V, G628S, L779P) were perfused with ICA-105574 (10 µM). The treatment induced a strong *I*_Kr_ activation on steady state currents in both WT and A561V (from 0 or -20 to +70 mV, respectively). In G628S, although the activation curve is not bell-shaped, ICA-105574 is able to induce a statistically significative difference (from -30 to +70 mV) partially restoring the functionality of the mutated channels. Notably, both the A561V and G628S mutations showed a statistically significant positive response to ICA-105574 (Figure 2 and S5) demonstrated by: 1) the increase of current amplitude in the I/V relationship (in A561V the peak is more than quadrupled in size); 2) the change of the shape of the I/V relationship, which exhibited a rectification, partially resembling the WT one (Figure 2). In A561V a complete bell-shaped curve was obtained, while in G628S the straight bent until to look like a half bell-shaped curve. This suggests that the *I*_Kr_ activator treatment could be able to induce an increment of the outward currents during the action potential plateau phase leading to acceleration of ventricular repolarization and a reduction in the duration of action potential and consequently shortening the QT phase. This is the first study demonstrating that A561V and G628S HERG mutations are responsive to ICA-105574 perfusion. Conversely, the HERG-L779P did not respond to ICA-105574, where *I*_Kr_ amplitude is not increased and I/V activation curve did not undergo any change of shape, suggesting that ICA-105574 does not induce any alteration in HERG protein’ conformation, preventing channel opening.

### Molecular Dynamics of WT, LQT2 and SQT1 HERG variants with and without ICA-105574

Having shown that ICA-105574 can rescue the activity of G628S and A561V *in vitro*, we set out to understand the molecular determinants of this rescue mechanism using MD simulations. We focused on the comparison of the ions permeability of the WT and G628S systems (Figure 3) alone or bound to ICA-105574 (WT-ICA, G628S-ICA). The G628S mutation, being in the SF, can have a direct effect on the conductivity of channel that is then rescued by ICA-105574. Of note, ICA-105574 binds relatively far from the mutation site. Furthermore, we simulated two SQT1 mutations, namely S631A, and N588K, to investigate whether the effect of ICA-105574 is structurally comparable to that of these pathological mutations, given their associated attenuated inactivation[9,10]. To demonstrate the effect of ICA-105574, two molecules were docked into the opposite subunit fenestrations based on the previously identified binding site residues [29–31] (Figure 3B). The rationale for using two ICA-105574 molecules is based on earlier study suggesting that a single ICA molecule binding to a subunit is insufficient to enhance channel activity [29]. ICA-105574 is found to be stable in WT-ICA and G628S-ICA systems over the course of simulations (Figure S7A, B). Our energetic analysis reveals that ICA-105574 molecules forms favorable polar and non-polar interactions in the form of H-bond (S649) and pi-pi interactions (F557, Y652, F656) in the respective fenestrations (Figure S7C, D).

**Figure 3.**
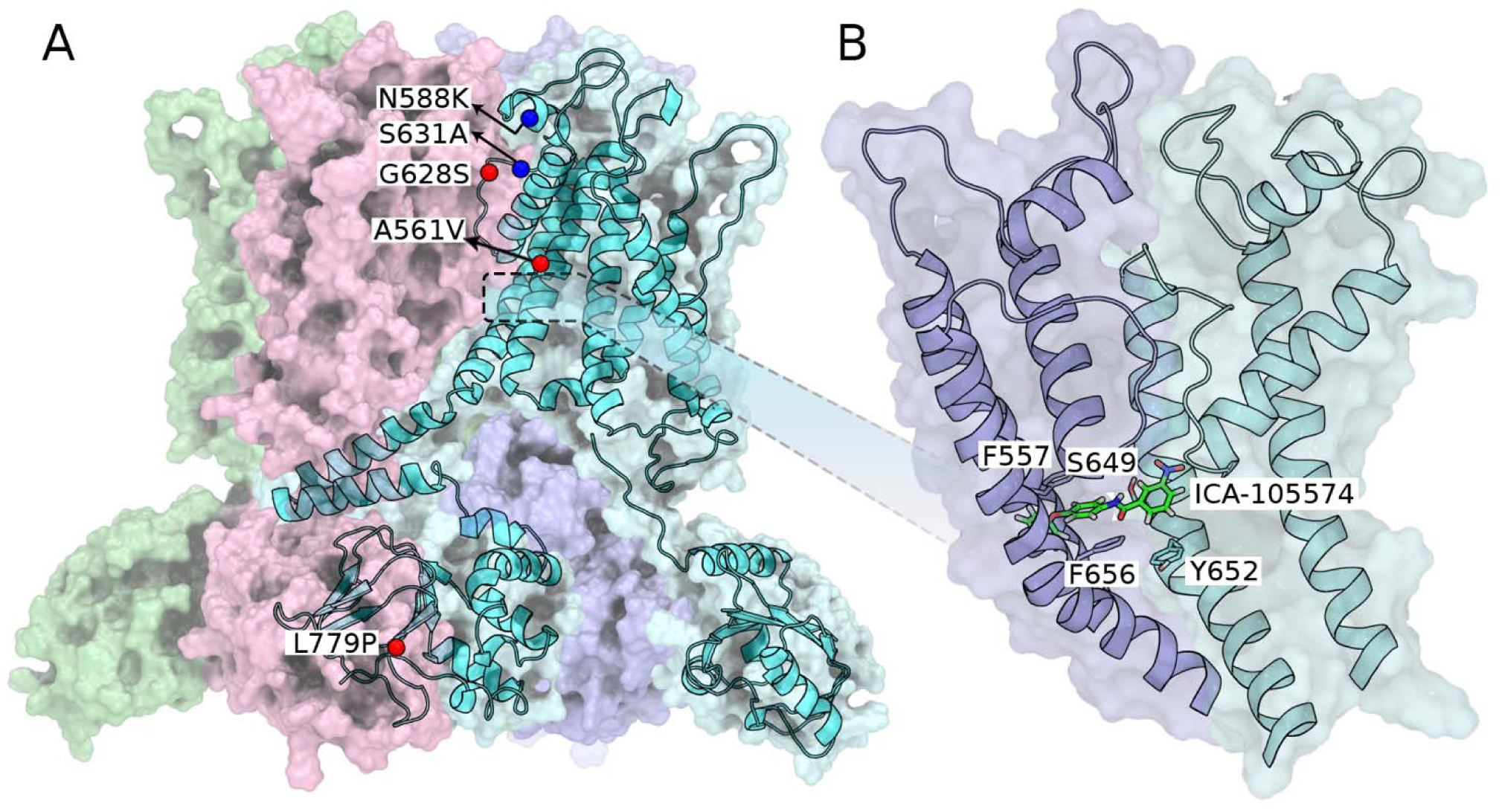
(A) Structural representation of the HERG tetramer with the SQT1 (N588K and S631A) and LQT2 (A561V, G628S and L779P) mutations highlighted in blue and red spheres respectively. A single monomer is shown with different domains identified in the cryo-EM structure (PDB: 5VA2). The SQT1 mutations, N588K and S631A, are located near the selectivity filter and are known to attenuate the inactivation state, whereas the LQT mutation G628S directly affects the selectivity filter. (B) ICA-105574 was docked into the HERG tetramer based on the previous mutagenesis study by Sanguinetti and co-workers (*28*–*30*). The docked structure is subjected to molecular dynamics (MD) simulations to obtain an equilibrated protein-ligand structure embedded in the lipid bilayer. The position of ICA-105574 in the equilibrated structure is consistent with previous findings and suggestions.

In a first round of preliminary simulations (three replicates performed at +350□mV and +700□mV each, 200□mM KCl, 310□K, ∼600-800□ns per run; Table S1A), replicating the work of Miranda et. al. [60], we observed, as previously reported, rapid inactivation on the ns time scale, manifested by the flipping of carbonyl oxygen atoms and subsequent SF distortion (Figure S6). In these simulations, very few ion permeation events were observed (except for the ions initially placed in the SF), followed by the intermittent collapse of the SF, which has been interpreted as a signature of the flickering ion current observed in HERG channels. Nonetheless, such a small number of transitions does not allow to obtain sufficient statistics to calculate the ionic conductance on the HERG active state. To overcome this limitation and ensure that the selectivity filter remains stable on the microsecond time scale, we restrained the backbone torsion angles (φ, ψ) for residues S624 to G628. These restraints have only the very localized effect of decreasing the likelihood of rotations of the backbone, without affecting other atomic motions including side-chains reorientations or modification of the size of the pore. This is done in the same spirit of symmetry restraints, or other restraints applied to compensate possible force-fields shortcomings [61,62]. Remarkably, simulations performed using the restraint (two replicates performed at ∼ +750□mV per system, 200□mM KCl, 310□K, 1–2□μs per run; Table S2) allowed us to record multiple K^+^ permeation events with an average transition time (τ) between 100 and 300 ns (Figure 4A) and corresponding to an ion conductance ranging from 0.82 ± 0.07 pS to 2.10 ± 0.06 pS across all protein variants, except for the severe G628S L mutation that do not show any conductance in agreement with our *in vitro* electrophysiology data (cf. Figure 2). Notably, in the wild-type simulations, we observed an ion conductance of 1.27 ± 0.04 pS, which increased to 1.67 ± 0.09 pS in the presence of the activator ICA-105574. Most importantly, the G628S-ICA system display K^+^ permeation (cf. Table S2) and an ion conductance of 1.72 ± 0.05 pS, in agreement with the ability of ICA-105574 to rescue such mutation *in vitro* (cf. Figure 2). Although the conductance values from our studies are underestimated by an order of magnitude compared to experimental data, which reports a single-channel conductance between 10 pS and 12.1 ± 0.8 [2,63,64], these discrepancies are consistent with those obtained for other potassium ion channels through in silico methods [65–68].

**Figure 4.**
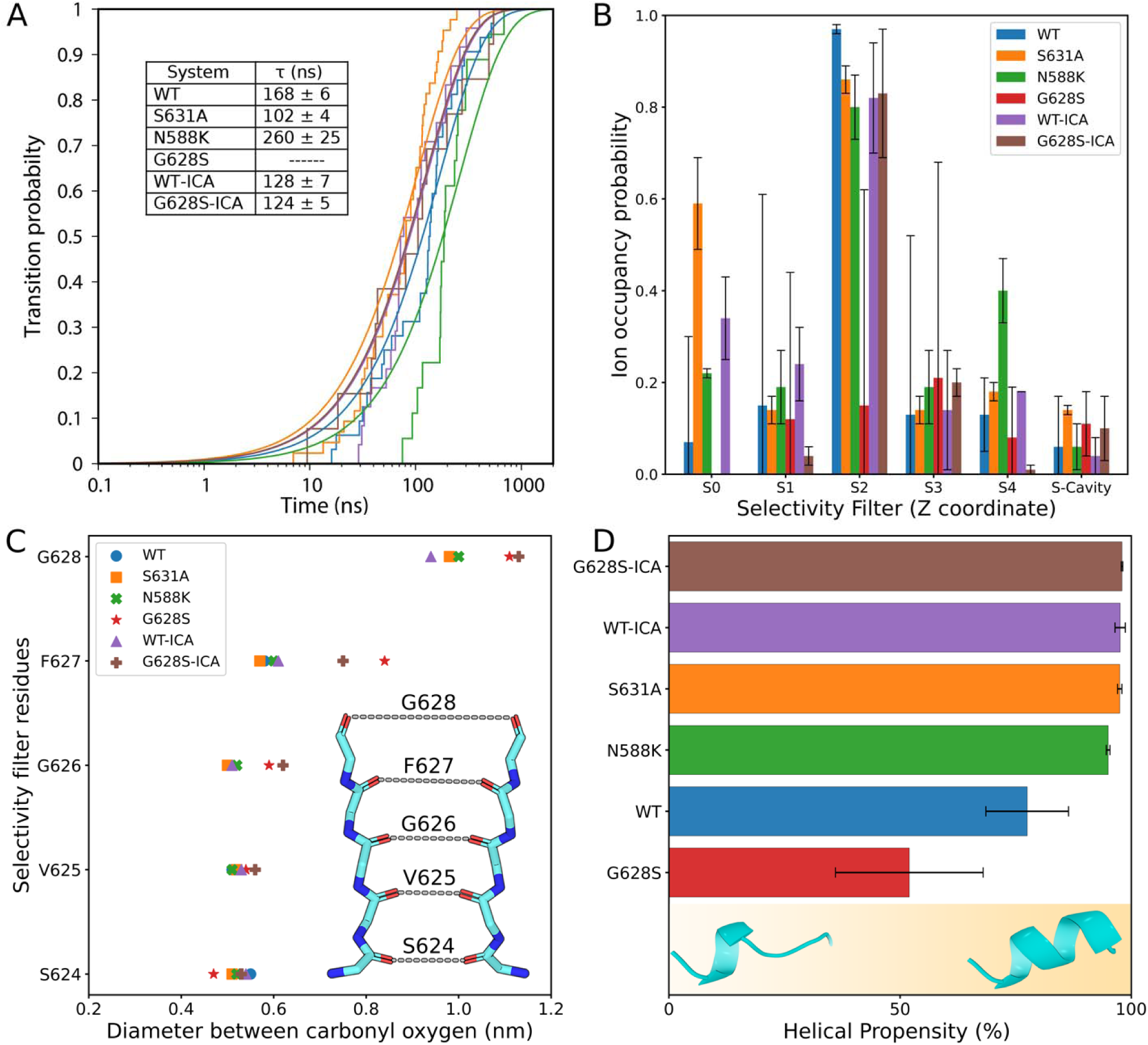
(A) The plot describes the transition probability of an ion crossing event and the average transition time for a single permeation event. No new ion permeation event was observed for the LQT2-G628S system except for the ions initially placed to prevent SF collapse. The inset table shows the average transition time for an ion permeation event in the HERG channel. (B) The plot represents the ion occupancy probability at different sites in the selectivity filter over the simulations. The selectivity filter is divided into six different sites namely S0 to S4 and S-Cavity. (C) Open state definition of SF: The plot shows the diameter of the SF averaged over all replica trajectories. The diameter is calculated by averaging over the interchain distance between the opposite carbonyl oxygens of the SF residues. The Mutation G628S increases the diameter of the SF at the extracellular side. (D) The plot shows the helical propensity (%) of the S5P turret helix (residue number 584 to 592) in different systems. The values were averaged over the monomers for the replicated trajectories. Two distinct population clusters were observed for the helical region. The lower helical propensity around 30% is a weakly ordered structure which is prominent in the G628S mutation, whereas the higher helical propensity (> 90%) is represented as a well-structured helix in WT, SQTs (N588K, S631A) and ICA-bound systems. For comparison, see Figure S9, which shows the population density of helical propensity in different systems.

Interestingly, our model effectively demonstrated the impact of ICA-105574 on the channel conductance in both the wild type and the G628S mutant. Altogether these results support the use of our restrained MD simulations to explore the effect of ICA-105574 and of mutations on the HERG active state.

### The helicity of the S5P turret as a signature of activation propensity

Having established a protocol that allowed us to accumulate statistics about HERG in its conducting state, we calculated the ion occupancy probability at different sites in the selectivity filter described previously for K^+^ ion channels (Figure 4B). All ion-permeable systems show higher ion occupancy at the S2 site, which in turn prevents the collapse of SF [69]. In contrast, the ion occupancy at S2 is drastically reduced in LQT2-G628S and is recovered in the presence of ICA-105574 in the G628S-ICA system. Structural and computational studies suggest that the active state of the HERG channel differs from inactivation in terms of the diameter of the SF and the heterogeneity in the dihedral angles of F627 [19,20,60]. In our simulations, the average diameter for the SF residues is found to be stable and similar for wild-type, SQT1 mutations (N588K, S631A), and WT-ICA systems (Figure 4C). While, in the presence of LQT2 mutation (G628S), there is a notable increase in the diameter around F627 (0.80 ± 0.1 nm) and S628 (1.13 ± 0.08 nm) showing the effect of the mutation. Of note, the G628S-ICA bound system showed only a small decrease in the SF diameter with respect to G628S, suggesting that in this range of values, SF diameter is not a clear signature of activation. To access any major conformational rearrangements and flexibility in the pore domain (PD) upon mutation or ligand binding, we analyzed the inter-residue pairwise distances for different systems with respect to the wild type (Figure S8 A-E). Overall, no major conformational changes in the pore domain were observed, except for a higher flexibility in the extracellular loops in the wild type and G628S as compared to the other systems (Figure S8F), suggesting that the differences we are looking for may be subtle and related to the conformational dynamics of the system. In HERG, experimental evidence emphasizes on the role of the S5P turret helix (residues 584-592) in the c-type inactivation mechanism [70–72], and the S5P segment has been observed to exist between helical and unstructured conformations in MD simulations. Interestingly, we observe a clear trend in the helicity of this region within SQT1 mutations and ligand-bound HERG channel as compared to the wild type (Figure 4D). Importantly, the G628S mutation has the least helical propensity, which is restored in the presence of ICA-105574. The activator is proposed to act primarily by attenuating rapid inactivation by increasing the stability of SF and channel conductance; it is intriguing how the binding of ICA-105574 below the SF can exert its function via conformational changes at long distance in the S5P turret helix. In contrast both SQT1 mutations are localized within or in contact with this region (Figure 3A) and may alter its role in c-type inactivation.

### ICA-105574 binding promotes the reorganization of hydrogen bonds through an allosteric network extending from the binding site to the S5P turret helix

Having observed that WT-ICA, G628S-ICA and the two SQT1 mutation share an increased helicity in the S5P turret helix in comparison to the WT and, even more, G628S systems, we investigated how ICA-105574 could exert such long-range structural effect. With this purpose we calculated the variation of the hydrogen bond (H-bond) occupancy in the pore domain of the HERG channel in the wild-type, G628S, and activator-bound G628S-ICA systems to identify changes in molecular interactions upon different perturbations (mutation or ligand binding). The net H-bond occupancy (%) between two residues i and j is given by 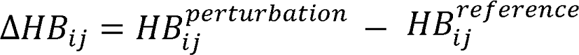 where the perturbation factor could be mutation or ligand binding while the reference could be either WT or G628S. As shown in Figure 5, red lines indicate residue pairs with higher H-bond occupancy (%) in the perturbed system, while blue lines indicate the same in the reference system (H-bond loss in perturbed system). When comparing G628S with WT, the analysis is the result of an averaging over the four chains (Figure 5A), whereas when comparing G628S with G628S-ICA, we had to take into account the asymmetric docking of the activator across two adjacent monomers, so we have analyzed the change in the H-bond occupancy in these chains separately (Figure 5B). Our analysis showed that the G628S mutation decreases the H-bond occupancy around the SF (F617-S621, S620-G626, S631-N633) and in the S5P turret helix spanning from G584 to Q592 as compared to the WT (Figure 5A). Notably, binding of ICA-105574 to the G628S system resulted in a marked increase in H-bond occupancy, in both adjacent chains, for residues around SF (Y616-S620, S620-G626) and in the S5P turret helix residues (G584-Q592).

**Figure 5.**
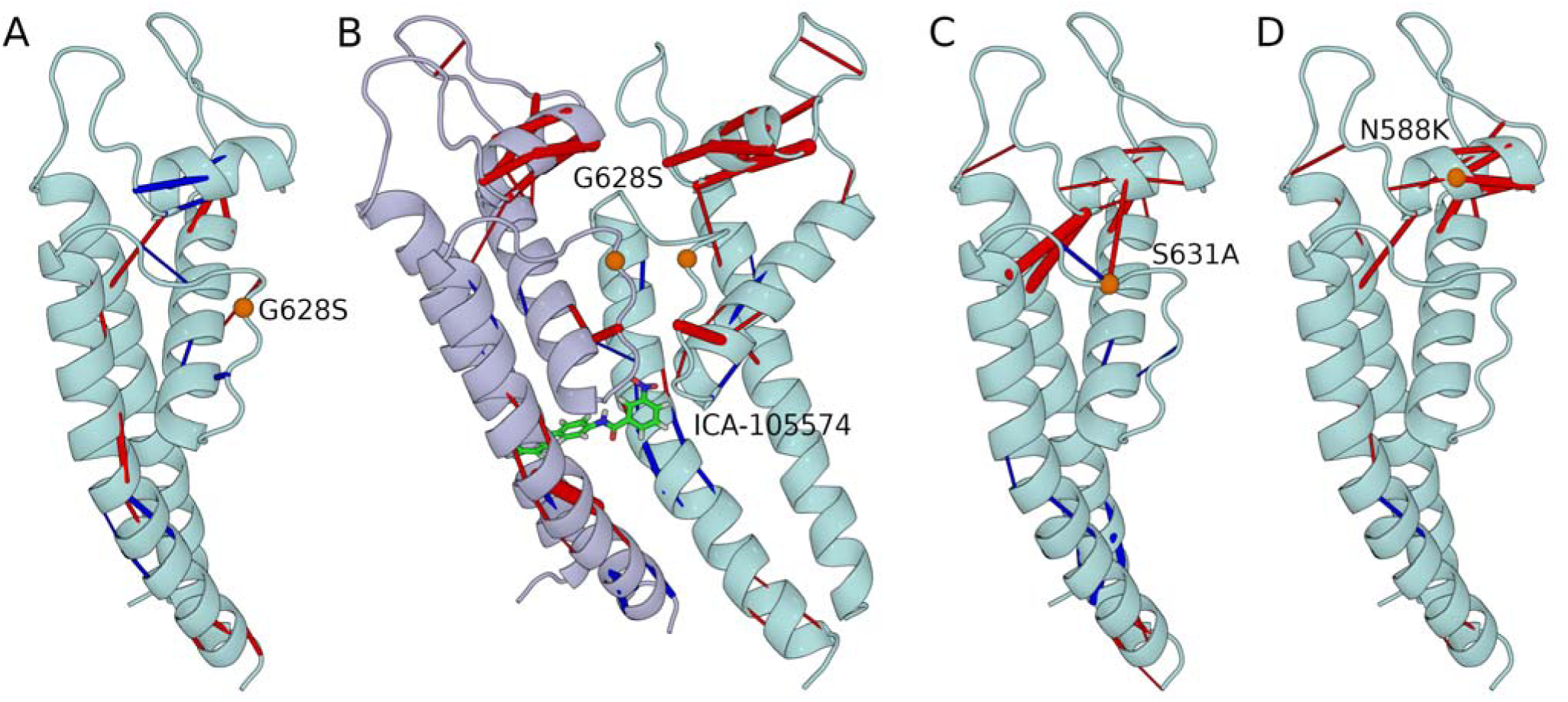
Hydrogen bond rearrangement upon perturbation in the form of mutations (G628S, S631A, N588K) or ligand binding (ICA-105574). The net H-bond occupancy (%) is highlighted in blue and red lines for the reference and the perturbed systems, respectively. The thickness of the lines represents the magnitude of the change. The mutations are shown as orange sphere. (A) Net H-bond occupancy (%) between G628S and WT 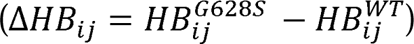 shows the effect of the mutation in wild-type HERG: Blue lines show H-bonds that are lost in the mutant around the SF and the S5P turret helix, while red lines show H-bonds that are gained in the mutant at some residue pairs and the S6 helix end. (B) Net H-bond occupancy (%) between G628S-ICA and G628S 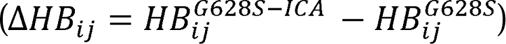 shows the rescue effect of ICA-105574: The figure shows two adjacent chains to represent different H-bonding patterns following the asymmetric binding of the activator in the HERG channel. ICA-105574 binding increases the H-bond occupancy around SF (Y616-S620, S620-G626) and S5P turret helix (G584-Q592) in both chains, while some residue pairs have opposite H-bond occupancy changes in G628S and ICA-105574 systems (e.g., E637-S641, M645-S649). (C, D) Net H-bond occupancy (%) between S631A and WT 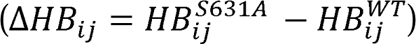, and between N588K and 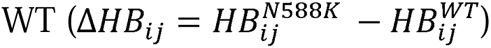 shows the effect of mutations in terms of gain in H-bonds in the mutants at S5P turret helix and some residue pairs around the S6 helix end.

The asymmetric positioning of ICA-105574 also resulted in different H-bonding patterns in adjacent chains. For example, one chain had higher H-bond occupancy near SF (F617-S621, S621-S624) and W568-W585, while another chain with more ICA-105574 contacts had increased H-bond occupancy in the S6 helical subunit. In addition, some residue pairs (e.g. Y616-N629, E637-S641, M645-S649 and at the S6 helix end) have higher H-bond occupancy in G628S than in WT (Figure 5A, red lines). However, this difference is reversed when ICA-105574 binds to G628S (Figure 5B, red lines). Notably, we also observed a similar H-bond rearrangement in the S5P turret region when comparing the WT and WT-ICA systems (Figure S10A), supporting that this is a robust feature of ICA-105574 binding. Furthermore, a similar pattern is observed when comparing the WT with the S631A and N588K SQT1variants (Figure 5 C, D). Overall, the increase in turret helicity (Figure 4D) and the decrease in RMSF (Figure S8F) observed above for both ICA-105574 bound and SQT1 systems can be attributed to a rearrangement of H-bonds occupancy, although the mechanism of stability may differ between systems. For example, the mutation of the polar amino acid (N588) to a charged amino acid (K588) increases helix stability by forming a salt bridge with D591, whereas mutation of S631A replaces the wild-type S631-N633 H-bond with an N588-A631 H-bond.

While these observations further highlight a similarity between the binding of ICA-105574 and two SQT1 mutations under study, they still do not fully describe how ICA-105574 may exert its long-range effect on the conformational dynamics of HERG. To this end, we focused on a single monomer chain and shortlisted residues with either higher net H-bond occupancy in G628S-ICA or *E_i,ICA_* < -2.5 *kcal/mol* (Figure S7D). We identified S624 as a common residue in this list and defined it as the starting point of the allosteric network, coupling ICA-105574 to the SF and the S5P turret helix. Specifically, we observed that the nitro group of ICA-105574, located behind the SF, formed favorable interaction with the hydroxyl group of S624 (Figure 6A). This resulted in a population shift of the chi (*χ*_1_) dihedral angle of S624, which describes the orientation of its side chain (Figure 6B). We hypothesize that this favorable interaction may trigger a domino effect in the form of an increase in the H-bond occupancy of several neighboring residue pairs behind the SF (Y616-S620, S620-G626), as previously shown in Figure 5, including A561-T618, F617-S621, S621-S624, with some of these occupancy changes manifesting as a significant shift in side chain orientation (e.g. T618, S620) upon ICA-105574 binding (Figure 6C). In particular, the changes observed for the region S620-S624 and the S5P turret helix are associated with a population shift in the *χ*-dihedral distribution of F617 and the tryptophan clamp (W568-W586).

**Figure 6.**
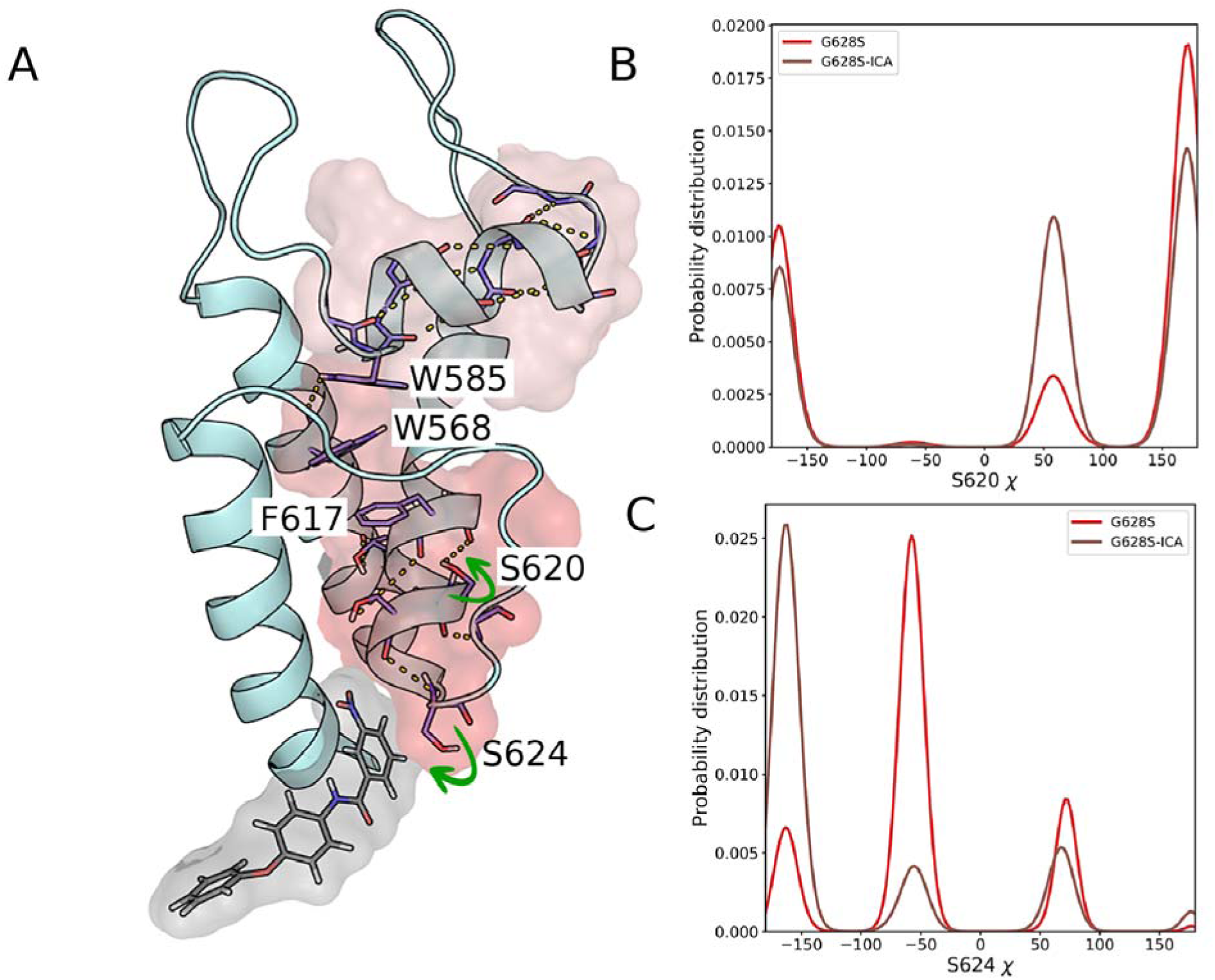
Mechanism of the rescue effect of the *I*_Kr_ activator ICA-105574 in the G628S mutation. (A) The figure shows a schematic representation of the allosteric network linking ICA-105574 binding to the SF and the S5P turret helix in a single monomer chain of the G628S mutant. ICA-105574 and the residues with higher net H-bond occupancy in the G628S-ICA system are highlighted as sticks, with H-bonds indicated by dashed lines. The surface representation shows the propagation path, with the gradual decrease in color indicating increasing distance from the ligand binding site. (B, C) S624 is defined as the starting point of the network because it forms a favorable interaction with the nitro group of ICA-105574. The Chi (χ)1 dihedral angle distributions of S620 and S624 are shown as 1D histograms, indicating significant population shifts or changes in the side chain orientations upon ICA-105574 binding.

In Figure 7, we have shown a 2D distribution of the *χ*_1_/ *χ*_2_ angle in the residue pairs W568-F617 (Figure 7A, B) and W568-W585 (Figure 7C, D). Here, we observed that the population density of the *χ* dihedrals for W568-F617 increases for a basin (F617(*χ*_2_) ∼ -100°, W568(*χ*_1_) ∼180°) without changing the overall shape of the distribution. While in the case of W568-W585, there is a marked shift for W585 (*χ*_1_) towards exclusively side chain orientation upon ICA-105574 binding, whereas the population for W568 (*χ*_1_) dihedral is significantly higher in one state (∼180°) in G628S-ICA compared to the G628S system. Thus, the pairing in the side chain orientations of these three residues may enhance the interaction between the S5, pore helix, and turret region in the activated state.

**Figure 7.**
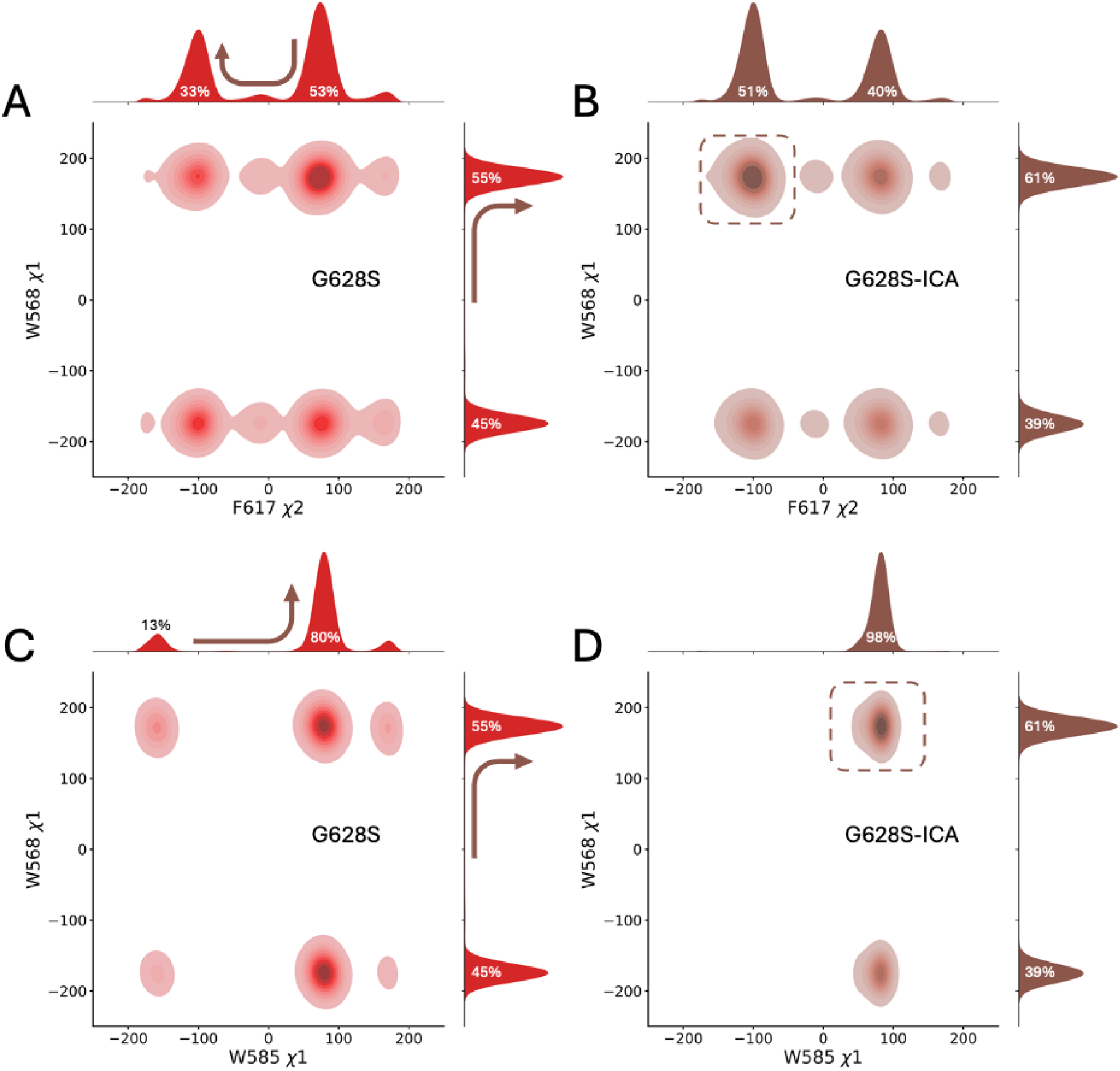
(A-D) *χ*2 dihedral distributions of residue pairs W568-F617 and W568-W585 in the G628S and G628S-ICA systems. The color opacity indicates the population density of each bin. The figure is plotted jointly for each axis variable with a 1D histogram to represent the population shift in the dihedrals. The arrows indicate the changes in the distribution upon ICA-105574 binding, while the dashed square box indicates the most populated basin in 2D distribution upon ICA-105574 binding. (A, B) show the 2D distribution of W568-*χ*1 and F617-*χ*2 angles, and (C, D) show the 2D distribution of *χ*1 angle for W585-W568.

Altogether these results highlight the strong interconnection among the residues in the vicinity of SF and how this connectivity may be exploited to stabilize it and rescue HERG activity. These results are also in agreement with the observation that mutation of any amino acid in this region is annotated as pathogenic or variant of concern, with essentially no benign mutations [73,74]. The A561V mutation also fall within this region. Of note, *in silico*, both the A561V and A561V-ICA systems exhibited ion permeation comparable to the WT and all conducting systems studied here, with ion occupancy in the SF consistent with the above observations (cf. Figure 4 and Figure S11). Interestingly, the ion conductance for A561V-ICA (i.e., 1.66 ± 0.12 pS) is nearly the same as that observed for the WT-ICA and G628S-ICA systems, indicating some robustness. No obvious differences were observed in the SF due to this mutation, furthermore S5P turret helicity is higher than WT (Figures S12, S13A). Additionally, the structural effect of ICA-105574 binding is comparable to that reported for G628S-ICA, with an increase of hydrogen bond occupancy extending from the binding site to the S5P turret helix (Figure S14). Also, the allosteric effect of ICA-105574 is robust, as evidenced by dihedral population shifts leading to conformational sidechain pairing between F617-W568 and W568-W585 in the A561V-ICA system, similar to those observed in G628S (Figure S15). Overall, these findings suggest that the A561V mutation does not impair conduction and may instead modulate the gating properties of HERG. Moreover, these results provide a mechanistic interpretation for the complex action of ICA-105574 on both gating and conductance. This is consistent with our *in vitro* electrophysiological data showing the different effect of ICA-105574 on G628S and A561V mutations, with the latter showing an altered steady state pre-pulse current amplitude and only partially distorted HERG current behavior.

In conclusion, our combined data allow to suggest that the A561V, G628S and L779P mutants act on different aspects of HERG function, with A561V possibly affecting gating but not conduction, G628S clearly disrupting the conductivity of the open state and L779P affecting protein localization as well function by strongly destabilizing the CNBHD domain (supported by additional CNBHD simulations, which showed increased fluctuations around the mutation, coupled with the loss of interactions in the inner core of the CNBHD; supplementary information and Figure S13B). Importantly, our data showed that the *I*_Kr_ activator ICA-105574 can rescue the A561V and G628S HERG LQT2 mutations to different extents by a mechanism that we propose is similar to that of some known SQT1 mutations (i.e., S631A and N588K). ICA-105574 binding stabilizes a network of amino acids in the SF, pore helix and S6 helix, allowing its effects to propagate ∼2 nm away to the S5P turret helix. Several residues (such as W568, W585, S621, T618) that are part of the identified allosteric pathway have been clinically or genetically implicated as pathogenic (involved in LQT or SQT) or of unknown clinical significance [73,74].

## Conclusions

LQTS predisposes young patients to develop ventricular fibrillation leading to sudden cardiac death. QT interval duration is a strong predictor of arrhythmic events, with each 10 ms increase conferring a 15% increased risk [75]. Priori and coworkers showed that substrate-specific therapy with mexiletine, which shortens the QT interval, confers arrhythmic protection in LQT3 [11,12]. Here we tested whether ICA-105574, an HERG activator, could be a good candidate to shorten the duration of cardiac repolarization in LQT2, by gaining an insight at three different levels: *in vivo*, *in vitro* and *in silico*.

*In vivo* experiments confirmed that administration of ICA-105574 induced a pronounced and statistically significant shortening of the QT interval in LQT2 and, importantly, no arrhythmic events were observed. In parallel, our *in vitro* and *in silico* analyses showed that ICA-105574 can partially rescue and restore the functionality of specific HERG LQT2 mutations by a promoting the reorganization of hydrogen bonds through an allosteric network extending from the ICA-105574 binding site to the S5P turret helix. We also propose that this mechanism is the same as that observed in gain-of-function SQT1 mutations. Thus, we support the hypothesis that HERG channel activators may represent a potentially very powerful novel therapeutic strategy for LQT2 patients. ICA-105574 by increasing the outward current may induce an acceleration of the delayed repolarizing phase of the ventricle of the LQT2 patients by shortening the QT interval.

Furthermore, we note that molecular dynamics simulations such as the one performed here, which focus on the functional effect of a drug are optimally positioned to provide an *in-silico* platform to test the functionality of HERG activators, even in the difficult case of testing a drug rescue mechanism.

## Supplementary materials

The additional supporting figures are available as separate file (Supporting Information). An illustration of the movement of potassium ions through the selectivity filter is shown in the form of a movie (mpg).

## Availability of data and materials

The MD simulations datasets generated and analyzed to support the conclusions of this article are available in Zenodo repository, https://zenodo.org/records/8349178.

## Ethics approval and consent to participate

The procedures employing Guinea pigs (Dunkin Hartley) were approved by the Ministero Italiano della Salute (Rome, Italy; 854/2019-PR).

## Author contributions

AK performed and analyzed molecular dynamics simulations; ET performed and analyzed in vitro experiments and performed the in vivo experiments; GL synthesized and developed the formulation for in vivo and in vitro delivery of the ICA-105574 compound; AT and DK performed and analyzed in vivo experiments; MD performed and analyzed the molecular biology experiments and performed the in vivo experiments; AK, ET, SGP, and CC wrote the manuscript with contributions from all authors; SGP and CC designed and supervised the project and provided resources. All authors have given approval to the final version of the manuscript.

## Acknowledgments

The authors thank Amanda Oldani, Patrizia Vaghi and Samantha Solito for technical assistance provided to the confocal microscopy and the cytofluorimetry facility, respectively, and “Centro Grandi Strumenti” of the University of Pavia. We also acknowledge a CINECA award under the ISCRA initiative (HP10C4059K), for the availability of high-performance computing resources and support and EuroHPC JU for awarding this project access to Lumi-C (EHPC-REG-2022R01-041) at CSC in Kajaani, Finland. We thank Gerhard Thiel (TU Darmstadt) for critical reading of the manuscript. The guinea pig and syringe images in Figure 1A were created using BioRender 2024 from BioRender.com. License Number: TR26ZK0LX7.

## References

1. Curran ME, Splawski I, Timothy KW, Vincen GM, Green ED, Keating MT. A molecular basis for cardiac arrhythmia: HERG mutations cause long QT syndrome. Cell [Internet]. 1995 [cited 2024 Feb 24];80:795–803. Available from: https://pubmed.ncbi.nlm.nih.gov/7889573/

2. Zou A, Curran ME, Keating MT, Sanguinetti MC. Single HERG delayed rectifier K+ channels expressed in Xenopus oocytes. American Journal of Physiology-Heart and Circulatory Physiology [Internet]. 1997;272:H1309–14. Available from: 10.1152/ajpheart.1997.272.3.H1309

3. Sanguinetti MC, Jiang C, Curran ME, Keating MT. A mechanistic link between an inherited and an acquird cardiac arrthytmia: HERG encodes the IKr potassium channel. Cell [Internet]. 1995 [cited 2024 Feb 24];81:299–307. Available from: http://www.cell.com/article/0092867495903402/fulltext

4. Crotti L, Celano G, Dagradi F, Schwartz PJ. Congenital long QT syndrome. Orphanet J Rare Dis [Internet]. 2008 [cited 2024 Mar 5];3:1–16. Available from: https://ojrd.biomedcentral.com/articles/10.1186/1750-1172-3-18

5. Anderson CL, Delisle BP, Anson BD, Kilby JA, Will ML, Tester DJ, et al. Most LQT2 Mutations Reduce Kv11.1 (hERG) Current by a Class 2 (Trafficking-Deficient) Mechanism. Circulation [Internet]. 2006 [cited 2024 Feb 25];113:365–73. Available from: https://www.ahajournals.org/doi/abs/10.1161/CIRCULATIONAHA.105.570200

6. Smith JL, Anderson CL, Burgess DE, Elayi CS, January CT, Delisle BP. Molecular pathogenesis of long QT syndrome type 2. J Arrhythm [Internet]. 2016 [cited 2024 Mar 5];32:373–80. Available from: https://onlinelibrary.wiley.com/doi/full/10.1016/j.joa.2015.11.009

7. Lees-Miller JP, Duan Y, Teng GQ, Thorstad K, Duff HJ. Novel gain-of-function mechanism in K(+) channel-related long-QT syndrome: altered gating and selectivity in the HERG1 N629D mutant. Circ Res [Internet]. 2000 [cited 2024 Feb 25];86:507–13. Available from: https://pubmed.ncbi.nlm.nih.gov/10720411/

8. Zhou Z, Gong Q, Epstein ML, January CT. HERG Channel Dysfunction in Human Long QT Syndrome. Journal of Biological Chemistry [Internet]. 1998 [cited 2024 Feb 25];273:21061–6. Available from: http://www.jbc.org/article/S002192581848951X/fulltext

9. Butler A, Zhang Y, Stuart AG, Dempsey CE, Hancox JC. Action potential clamp characterization of the S631A hERG mutation associated with short QT syndrome. Physiol Rep [Internet]. 2018 [cited 2024 Feb 23];6. Available from: /pmc/articles/PMC6119704/

10. McPate MJ, Duncan RS, Milnes JT, Witchel HJ, Hancox JC. The N588K-HERG K+ channel mutation in the ‘short QT syndrome’: Mechanism of gain-in-function determined at 37 °C. Biochem Biophys Res Commun. 2005;334:441–9.

11. Priori SG, Napolitano C, Cantù F, Brown AM, Schwartz PJ. Differential response to Na+ channel blockade, beta-adrenergic stimulation, and rapid pacing in a cellular model mimicking the SCN5A and HERG defects present in the long-QT syndrome. Circ Res [Internet]. 1996 [cited 2024 Feb 22];78:1009–15. Available from: https://pubmed.ncbi.nlm.nih.gov/8635231/

12. Mazzanti A, Maragna R, Faragli A, Monteforte N, Bloise R, Memmi M, et al. Gene-Specific Therapy With Mexiletine Reduces Arrhythmic Events in Patients With Long QT Syndrome Type 3. J Am Coll Cardiol [Internet]. 2016 [cited 2024 Feb 22];67:1053–8. Available from: https://pubmed.ncbi.nlm.nih.gov/26940925/

13. Sanguinetti MC. HERG1 Channel Agonists and Cardiac Arrhythmia. Curr Opin Pharmacol [Internet]. 2014 [cited 2024 Feb 22];0:22. Available from: /pmc/articles/PMC3984452/

14. Vandenberg JI, Perry MD, Perrin MJ, Mann SA, Ke Y, Hill AP, et al. hERG K CHANNELS: STRUCTURE, FUNCTION, AND CLINICAL SIGNIFICANCE. Physiol Rev [Internet]. 2012;92:1393–478. Available from: www.prv.org

15. Li J, Shen R, Reddy B, Perozo E, Roux B. Mechanism of C-type inactivation in the hERG potassium channel. Sci Adv [Internet]. 2021 [cited 2024 Feb 22];7. Available from: https://pubmed.ncbi.nlm.nih.gov/33514547/

16. Pettini F, Domene C, Furini S. Early Steps in C-Type Inactivation of the hERG Potassium Channel. J Chem Inf Model. 2023;63:251–8.

17. Costa F, Guardiani C, Giacomello A. Molecular dynamics simulations suggest possible activation and deactivation pathways in the hERG channel. Communications Biology 2022 5:1 [Internet]. 2022 [cited 2024 Feb 25];5:1–11. Available from: https://www.nature.com/articles/s42003-022-03074-9

18. Wang W, MacKinnon R. Cryo-EM Structure of the Open Human Ether-à-go-go-Related K+ Channel hERG. Cell. 2017;169:422–430.e10.

19. Ngo K, Yarov-Yarovoy V, Clancy CE, Vorobyov I. Harnessing AlphaFold to reveal state secrets: Prediction of hERG closed and inactivated states. bioRxiv [Internet]. 2024 [cited 2024 Feb 26];2024.01.27.577468. Available from: https://www.biorxiv.org/content/10.1101/2024.01.27.577468v1

20. Lau CHY, Flood E, Hunter MJ, Williams-Noonan BJ, Corbett KM, Ng C-A, et al. Potassium dependent structural changes in the selectivity filter of HERG potassium channels. Nat Commun [Internet]. 2024;15:7470. Available from: 10.1038/s41467-024-51208-w

21. Asai T, Adachi N, Moriya T, Oki H, Maru T, Kawasaki M, et al. Cryo-EM Structure of K+-Bound hERG Channel Complexed with the Blocker Astemizole. Structure. 2021;29:203–212.e4.

22. Ekins S, Crumb WJ, Dustan Sarazan R, Wikel JH, Wrighton SA. Three-Dimensional Quantitative Structure-Activity Relationship for Inhibition of Human Ether-a-Go-Go-Related Gene Potassium Channel. Journal of Pharmacology and Experimental Therapeutics [Internet]. 2002 [cited 2024 Feb 25];301:427–34. Available from: https://jpet.aspetjournals.org/content/301/2/427

23. Creanza TM, Delre P, Ancona N, Lentini G, Saviano M, Mangiatordi GF. Structure-Based Prediction of hERG-Related Cardiotoxicity: A Benchmark Study. J Chem Inf Model [Internet]. 2021 [cited 2024 Feb 22];61:4758–70. Available from: https://pubs.acs.org/doi/full/10.1021/acs.jcim.1c00744

24. Cavalli A, Poluzzi E, De Ponti F, Recanatini M. Toward a pharmacophore for drugs inducing the long QT syndrome: Insights from a CoMFA study of HERG K+ channel blockers. J Med Chem [Internet]. 2002 [cited 2024 Feb 22];45:3844–53. Available from: https://pubs.acs.org/doi/abs/10.1021/jm0208875

25. Zhou PZ, Babcock J, Liu LQ, Li M, Gao ZB. Activation of human ether-a-go-go related gene (hERG) potassium channels by small molecules. Acta Pharmacol Sin. 2011. p. 781– 8.

26. El Harchi A, Brincourt O. Pharmacological activation of the hERG K+ channel for the management of the long QT syndrome: A review. J Arrhythm [Internet]. 2022 [cited 2024 Feb 23];38:554–69. Available from: https://pubmed.ncbi.nlm.nih.gov/35936037/

27. Asayama M, Kurokawa J, Shirakawa K, Okuyama H, Kagawa T, Okada JI, et al. Effects of an hERG activator, ICA-105574, on electrophysiological properties of canine hearts. J Pharmacol Sci. 2013;121:1–8.

28. Gerlach AC, Stoehr SJ, Castle NA. Pharmacological Removal of Human Ether-à-go-go-Related Gene Potassium Channel Inactivation by 3-Nitro-N-(4-phenoxyphenyl) Benzamide (ICA-105574). Mol Pharmacol [Internet]. 2010 [cited 2024 Feb 26];77:58–68. Available from: https://molpharm.aspetjournals.org/content/77/1/58

29. Zangerl-Plessl EM, Berger M, Drescher M, Chen Y, Wu W, Maulide N, et al. Toward a Structural View of hERG Activation by the Small-Molecule Activator ICA-105574. J Chem Inf Model. 2020;60:360–71.

30. Garg V, Stary-Weinzinger A, Sanguinetti MC. ICA-105574 Interacts with a Common Binding Site to Elicit Opposite Effects on Inactivation Gating of EAG and ERG Potassium Channels. Mol Pharmacol [Internet]. 2013 [cited 2024 Feb 25];83:805–13. Available from: https://molpharm.aspetjournals.org/content/83/4/805

31. Garg V, Stary-Weinzinger A, Sachse F, Sanguinetti MC. Molecular Determinants for Activation of Human Ether-à-go-go-related Gene 1 Potassium Channels by 3-Nitro-N-(4-phenoxyphenyl) Benzamide. Mol Pharmacol [Internet]. 2011 [cited 2024 Feb 25];80:630–7. Available from: https://molpharm.aspetjournals.org/content/80/4/630

32. Wu W, Sachse FB, Gardner A, Sanguinetti MC. Stoichiometry of altered hERG1 channel gating by small molecule activators. Journal of General Physiology [Internet]. 2014 [cited 2024 Feb 25];143:499–512. Available from: www.jgp.org/cgi/doi/10.1085/jgp.201311038

33. Porta-Sánchez A, Mazzanti A, Tarifa C, Kukavica D, Trancuccio A, Mohsin M, et al. Unexpected impairment of INa underpins reentrant arrhythmias in a knock-in swine model of Timothy syndrome. Nature Cardiovascular Research 2023 2:12 [Internet]. 2023 [cited 2024 Mar 6];2:1291–309. Available from: https://www.nature.com/articles/s44161-023-00393-w

34. Ruppert S, Vormberge T, Igl BW, Hoffmann M. ECG telemetry in conscious guinea pigs. J Pharmacol Toxicol Methods [Internet]. 2016 [cited 2024 Feb 24];81:88–98. Available from: https://pubmed.ncbi.nlm.nih.gov/27118261/

35. Wood AJJ, Hohnloser SH, Woosley RL. Sotalol. 101056/NEJM199407073310108 [Internet]. 1994 [cited 2024 Mar 7];331:31–8. Available from: https://www.nejm.org/doi/full/10.1056/NEJM199407073310108

36. Gałecki A, Burzykowski T. Linear Mixed-Effects Models Using R. 2013 [cited 2024 Mar 7]; Available from: https://link.springer.com/10.1007/978-1-4614-3900-4

37. Kenward MG, Roger JH. Small sample inference for fixed effects from restricted maximum likelihood. Biometrics [Internet]. 1997 [cited 2024 Mar 7];53:983. Available from: https://pubmed.ncbi.nlm.nih.gov/9333350/

38. Milani G, Budriesi R, Tavazzani E, Cavalluzzi MM, Mattioli LB, Miniero DV, et al. hERG stereoselective modulation by mexiletine-derived ureas: Molecular docking study, synthesis, and biological evaluation. Arch Pharm (Weinheim). 2023;

39. Friesner RA, Banks JL, Murphy RB, Halgren TA, Klicic JJ, Mainz DT, et al. Glide: A New Approach for Rapid, Accurate Docking and Scoring. 1. Method and Assessment of Docking Accuracy. J Med Chem [Internet]. 2004 [cited 2024 Mar 6];47:1739–49. Available from: https://pubs.acs.org/doi/abs/10.1021/jm0306430

40. Jo S, Kim T, Iyer VG, Im W. CHARMM-GUI: A web-based graphical user interface for CHARMM. J Comput Chem [Internet]. 2008 [cited 2024 Mar 7];29:1859–65. Available from: https://onlinelibrary.wiley.com/doi/full/10.1002/jcc.20945

41. Eswar N, John B, Mirkovic N, Fiser A, Ilyin VA, Pieper U, et al. Tools for comparative protein structure modeling and analysis. Nucleic Acids Res [Internet]. 2003 [cited 2024 Mar 7];31:3375–80. Available from: https://pubmed.ncbi.nlm.nih.gov/12824331/

42. Jorgensen WL, Chandrasekhar J, Madura JD, Impey RW, Klein ML. Comparison of simple potential functions for simulating liquid water. J Chem Phys [Internet]. 1983 [cited 2024 Mar 6];79:926–35. Available from: /aip/jcp/article/79/2/926/776316/Comparison-of-simple-potential-functions-for

43. Huang J, Rauscher S, Nawrocki G, Ran T, Feig M, De Groot BL, et al. CHARMM36m: An Improved Force Field for Folded and Intrinsically Disordered Proteins. Nat Methods [Internet]. 2017 [cited 2024 Mar 6];14:71. Available from: /pmc/articles/PMC5199616/

44. Noskov SY, Bernéche S, Roux B. Control of ion selectivity in potassium channels by electrostatic and dynamic properties of carbonyl ligands. Nature 2004 431:7010 [Internet]. 2004 [cited 2024 Feb 27];431:830–4. Available from: https://www.nature.com/articles/nature02943

45. Vanommeslaeghe K, MacKerell AD. Automation of the CHARMM general force field (CGenFF) I: Bond perception and atom typing. J Chem Inf Model [Internet]. 2012 [cited 2024 Mar 7];52:3144–54. Available from: https://pubs.acs.org/doi/abs/10.1021/ci300363c

46. Tribello GA, Bonomi M, Branduardi D, Camilloni C, Bussi G. PLUMED 2: New feathers for an old bird. Comput Phys Commun. 2014;185:604–13.

47. Bussi G, Donadio D, Parrinello M. Canonical sampling through velocity rescaling. Journal of Chemical Physics [Internet]. 2007 [cited 2024 Mar 7];126. Available from: /aip/jcp/article/126/1/014101/186581/Canonical-sampling-through-velocity-rescaling

48. Bernetti M, Bussi G. Pressure control using stochastic cell rescaling. Journal of Chemical Physics [Internet]. 2020 [cited 2024 Mar 7];153. Available from: /aip/jcp/article/153/11/114107/199610/Pressure-control-using-stochastic-cell-rescaling

49. Darden T, York D, Pedersen L. Particle mesh Ewald: An N⋅log(N) method for Ewald sums in large systems. J Chem Phys [Internet]. 1993 [cited 2024 Mar 7];98:10089–92. Available from: /aip/jcp/article/98/12/10089/461765/Particle-mesh-Ewald-An-N-log-N-method-for-Ewald

50. Hess B, Bekker H, Berendsen HJC, Fraaije JGEM. LINCS: A linear constraint solver for molecular simulations. J Comput Chem [Internet]. 1997;18:1463–72. Available from: 10.1002/(SICI)1096-987X(199709)18:12<1463::AID-JCC4>3.0.CO

51. Abraham MJ, Murtola T, Schulz R, Páll S, Smith JC, Hess B, et al. GROMACS: High performance molecular simulations through multi-level parallelism from laptops to supercomputers. SoftwareX. 2015;1–2:19–25.

52. Li Y, Ng HQ, Li Q, Kang CB. Structure of the Cyclic Nucleotide-Binding Homology Domain of the hERG Channel and Its Insight into Type 2 Long QT Syndrome. Scientific Reports 2016 6:1 [Internet]. 2016 [cited 2024 Mar 7];6:1–10. Available from: https://www.nature.com/articles/srep23712

53. Sakaguchi Y, Sugiyama A, Takao S, Akie Y, Takahara A, Hashimoto K. Halothane Sensitizes the Guinea-Pig Heart to Pharmacological I_Kr_ Blockade: Comparison With Urethane Anesthesia. J Pharmacol Sci. 2005;99:185–90.

54. Takahara A, Sugiyama A, Hashimoto K. Reduction of repolarization reserve by halothane anaesthesia sensitizes the guinea-pig heart for drug-induced QT interval prolongation. Br J Pharmacol [Internet]. 2005;146:561–7. Available from: 10.1038/sj.bjp.0706352

55. Busch AE, Malloy K, Groh WJ, Varnum MD, Adelman JP, Maylie J. The Novel Class III Antiarrhythmics NE-10064 and NE-10133 Inhibit Isk Channels Expressed in Xenopus Oocytes and Iks in Guinea Pig Cardiac Myocytes. Biochem Biophys Res Commun [Internet]. 1994;202:265–70. Available from: https://www.sciencedirect.com/science/article/pii/S0006291X8471922X

56. Numaguchi H, Mullins FM, Johnson JP, Johns DC, Po SS, Yang ICH, et al. Probing the interaction between inactivation gating and Dd-sotalol block of HERG. Circ Res [Internet]. 2000 [cited 2024 Feb 22];87:1012–8. Available from: https://pubmed.ncbi.nlm.nih.gov/11090546/

57. Rasmussen HS, Allen MJ, Blackburn KJ, Butrous GS, Dalrymple HW. Dofetilide, a novel class III antiarrhythmic agent. J Cardiovasc Pharmacol [Internet]. 1992 [cited 2024 Feb 24];20 Suppl 2:S96–105. Available from: https://pubmed.ncbi.nlm.nih.gov/1279316/

58. Sanguinetti MC, Curran ME, Spector PS, Keating MT. Spectrum of HERG K+-channel dysfunction in an inherited cardiac arrhythmia. Proceedings of the National Academy of Sciences [Internet]. 1996 [cited 2024 Feb 25];93:2208–12. Available from: https://www.pnas.org/doi/abs/10.1073/pnas.93.5.2208

59. Stump MR, Gong Q, Zhou Z. Isoform-Specific Dominant-Negative Effects Associated with hERG1 G628S Mutation in Long QT Syndrome. PLoS One [Internet]. 2012 [cited 2024 Feb 25];7:e42552. Available from: https://journals.plos.org/plosone/article?id=10.1371/journal.pone.0042552

60. Miranda WE, DeMarco KR, Guo J, Duff HJ, Vorobyov I, Clancy CE, et al. Selectivity filter modalities and rapid inactivation of the hERG1 channel. Proc Natl Acad Sci U S A [Internet]. 2020 [cited 2024 Feb 22];117:2795–804. Available from: https://www.pnas.org/doi/abs/10.1073/pnas.1909196117

61. Porro A, Saponaro A, Gasparri F, Bauer D, Gross C, Pisoni M, et al. The HCN domain couples voltage gating and cAMP response in hyperpolarization-activated cyclic nucleotide-gated channels. Boudker O, Csanády L, Csanády L, Bankston J, Goldschen-Ohm MP, editors. Elife [Internet]. 2019;8:e49672. Available from: 10.7554/eLife.49672

62. Camilloni C, Vendruscolo M. Statistical Mechanics of the Denatured State of a Protein Using Replica-Averaged Metadynamics. J Am Chem Soc [Internet]. 2014;136:8982–91. Available from: 10.1021/ja5027584

63. Kiehn J, Lacerda AE, Wible B, Brown AM. Molecular Physiology and Pharmacology of HERG. Circulation [Internet]. 1996;94:2572–9. Available from: 10.1161/01.CIR.94.10.2572

64. Veldkamp MW, van Ginneken ACG, Opthof T, Bouman LN. Delayed Rectifier Channels in Human Ventricular Myocytes. Circulation [Internet]. 1995;92:3497–504. Available from: 10.1161/01.CIR.92.12.3497

65. Furini S, Domene C. Critical Assessment of Common Force Fields for Molecular Dynamics Simulations of Potassium Channels. J Chem Theory Comput [Internet]. 2020;16:7148–59. Available from: 10.1021/acs.jctc.0c00331

66. Lam CK, de Groot BL. Ion Conduction Mechanisms in Potassium Channels Revealed by Permeation Cycles. J Chem Theory Comput [Internet]. 2023;19:2574–89. Available from: 10.1021/acs.jctc.3c00061

67. Jäger M, Koslowski T, Wolf S. Predicting Ion Channel Conductance via Dissipation-Corrected Targeted Molecular Dynamics and Langevin Equation Simulations. J Chem Theory Comput [Internet]. 2022;18:494–502. Available from: 10.1021/acs.jctc.1c00426

68. Ocello R, Furini S, Lugli F, Recanatini M, Domene C, Masetti M. Conduction and Gating Properties of the TRAAK Channel from Molecular Dynamics Simulations with Different Force Fields. J Chem Inf Model [Internet]. 2020;60:6532–43. Available from: 10.1021/acs.jcim.0c01179

69. Boiteux C, Posson DJ, Allen TW, Nimigean CM. Selectivity filter ion binding affinity determines inactivation in a potassium channel. Proc Natl Acad Sci U S A [Internet]. 2020 [cited 2024 Feb 26];117:29968–78. Available from: https://www.pnas.org/doi/abs/10.1073/pnas.2009624117

70. Liu J, Zhang M, Jiang M, Tseng GN. Structural and functional role of the extracellular S5-P linker in the HERG potassium channel. Journal of General Physiology. 2002;120:723–37.

71. Torres AM, Bansal PS, Sunde M, Clarke CE, Bursill JA, Smith DJ, et al. Structure of the HERG K+ Channel S5P Extracellular Linker: ROLE OF AN AMPHIPATHIC α-HELIX IN C-TYPE INACTIVATION. Journal of Biological Chemistry. 2003;278:42136–48.

72. Clarke CE, Hill AP, Zhao J, Kondo M, Subbiah RN, Campbell TJ, et al. Effect of S5P α-helix charge mutants on inactivation of hERG K+ channels. J Physiol [Internet]. 2006 [cited 2024 Feb 27];573:291. Available from: /pmc/articles/PMC1779719/

73. Anderson CL, Kuzmicki CE, Childs RR, Hintz CJ, Delisle BP, January CT. Large-scale mutational analysis of Kv11.1 reveals molecular insights into type 2 long QT syndrome. Nat Commun. 2014;5.

74. Kapplinger JD, Tester DJ, Salisbury BA, Carr JL, Harris-Kerr C, Pollevick GD, et al. Spectrum and prevalence of mutations from the first 2,500 consecutive unrelated patients referred for the FAMILION® long QT syndrome genetic test. Heart Rhythm. 2009;6:1297–303.

75. Mazzanti A, Maragna R, Vacanti G, Monteforte N, Bloise R, Marino M, et al. Interplay Between Genetic Substrate, QTc Duration, and Arrhythmia Risk in Patients With Long QT Syndrome. J Am Coll Cardiol. 2018;71:1663–71.

