## Supporting Information for "Molecular Insights into the Rescue Mechanism of an HERG Activator Against Severe LQT2 Mutations"

**This PDF file includes:**

Supplementary Methods

Supplementary Results

Supplementary Figures S1 to S14

Supplementary Tables S1 to S6

Legends for Movies S1

Legends for Datasets S1

**Other supplementary materials for this manuscript include the following:**

Movies S1

### Supplementary Methods

#### Cloning of hERG-WT and mutants in heterologous systems

Primers used for mutagenesis:

A561V-For-CTTTGCGCTCATCGtGCACTGGCTAGCCT,

A561V-Rev-AGGCTAGCCAGTGCACGATGAGCGCAAAG,

G628S-For-GGGAGAGACGTTGCTGAAGCCCACACTGG,

G628S-Rev-CCAGTGTGGGCTTCAGCAACGTCTCTCCC,

L779P-For-GGGAGATGAAGTACGGGGCGGTGAGCAGG,

L779P-Rev-CCTGCTCACCGCCCCGTACTTCATCTCCC

Primers used for subcloning in pmCherry-N1:

Subcloning Xho I_For-TTGCCTCGAGGAGTACTTCCAG

Subcloning Xho I_Rev-AAACCTCGAGGCGCTGGCGCAGG

Age I_YFP_for-ACACCGGTCGCCACCATGGTGAG

Not I_YFP_Rev-ACTCGGCCGCTACTTGTACAGCTCGTCCAT

### Supplementary Results

#### Molecular Dynamics simulations of the L779P LQT2 variant

For the the L779P mutation, located in the cyclic nucleotide-binding homology domain (CNBHD), we focused on simulating only the CNBHD of HERG channels [1] in both the wild-type and mutant states. The L779 residue is strategically positioned within the central beta-strand of the beta-roll, which is a common fold among canonical cyclic nucleotide-binding proteins. It establishes favorable hydrophobic and backbone H-bond interactions with neighboring residues, thus contributing to the overall stability of this domain. However, the introduction of a proline mutation at position L779 can have a significant impact on the overall structure of the CNBHD. Interestingly, our simulations showed a significant increase in the RMSF for the CNBHD in the presence of the L779P mutation (Figure S11B). The dramatic increase in fluctuations observed here, coupled with the loss of interactions in the inner core of the CNBHD, suggests a loss of stability in this domain, potentially increasing the risk of misfolding and a putative loss of functionality. This is consistent with our *in vitro* observations, where most of the protein is localized in the cytoplasm, with enrichment around the perinuclear region, suggesting some folding and trafficking defects. Having observed that the effect of ICA-105574 is primarily localized to the SF region, its inability to rescue such a mutation is therefore expected.

### Supplementary figures

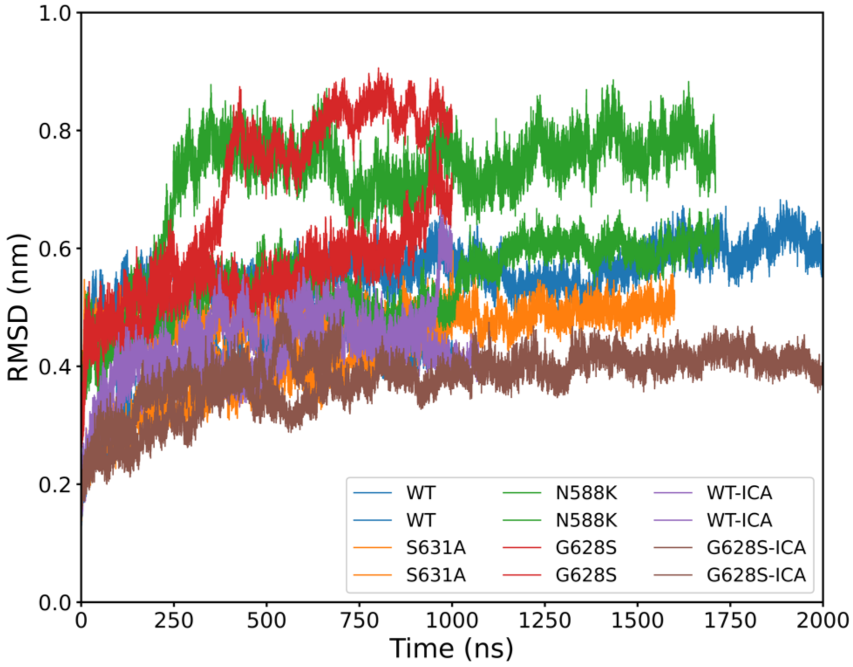

**Figure S1.** Backbone root mean square deviation (RMSD) comparison with minimized structures for each system. All systems remained stable and well equilibrated throughout the production runs. Notable deviations in the RMSD of specific replicas of the N588K system and G628S-ICA were further analyzed by calculating the RMSD for the pore domain of the individual trajectories.

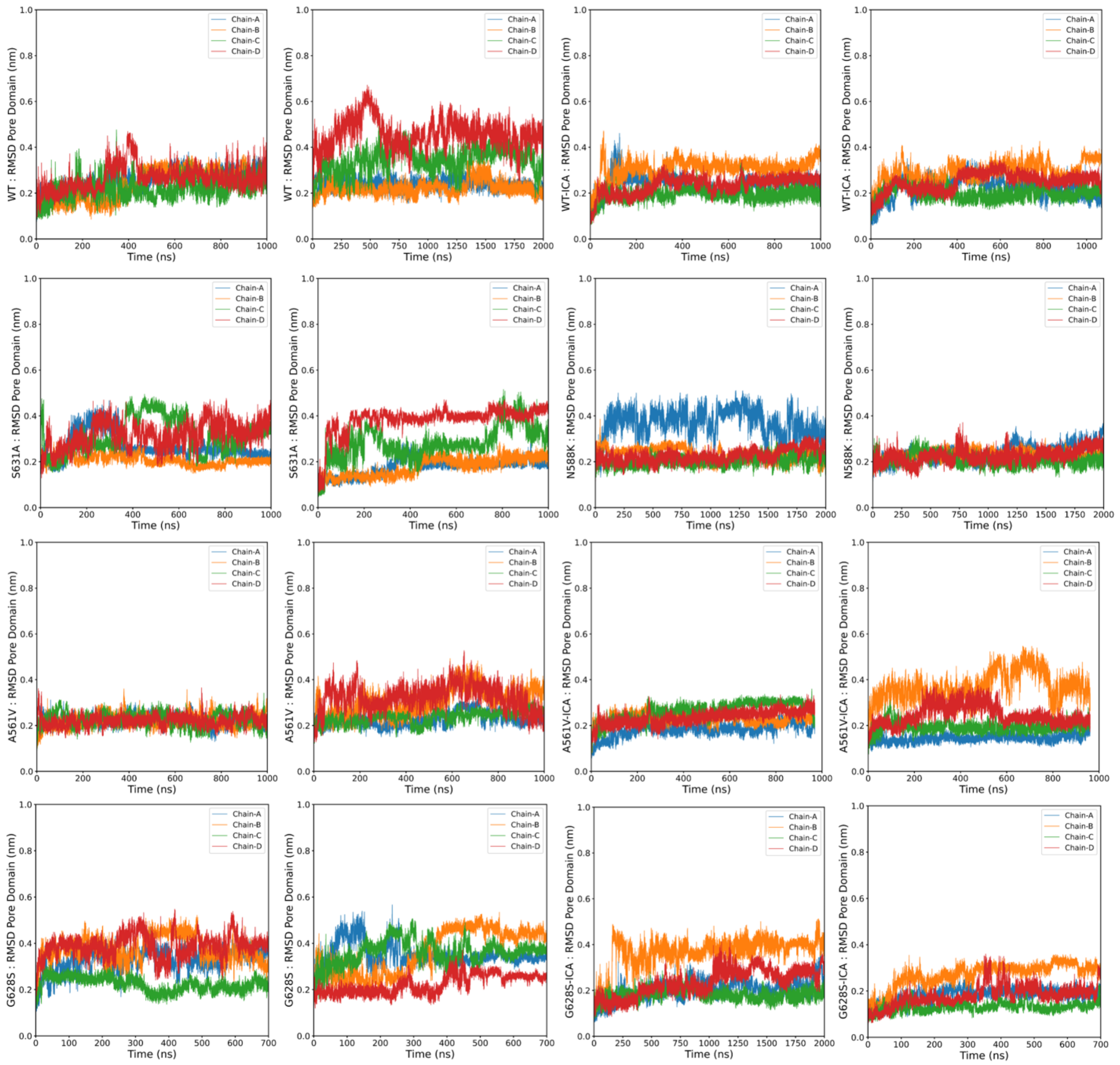

**Figure S2.** RMSD for the pore domain of individual monomers across multiple replicas for different systems.

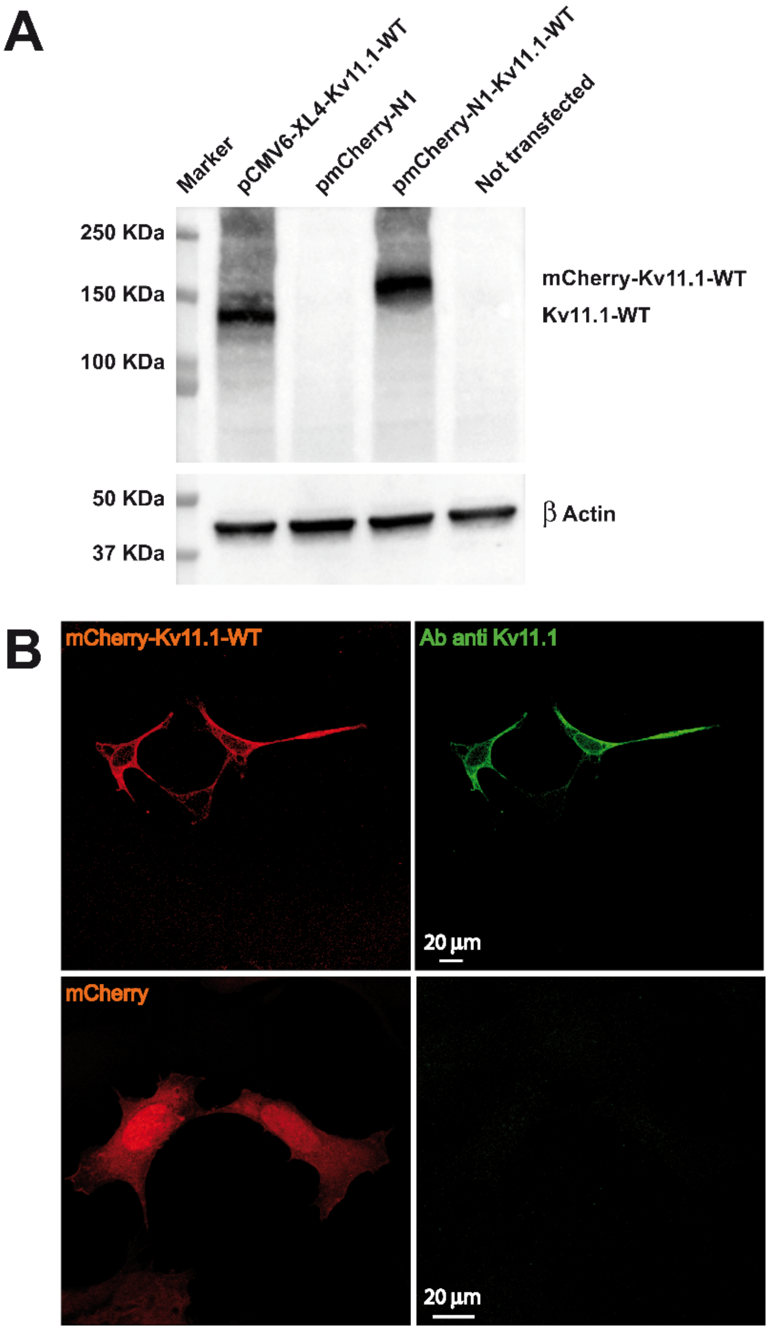

**Figure S3.** Analysis of protein expression of hERG-WT after transient transfection in HEK293 cells. (A) Western blot demonstrates the correct migration and expected molecular weight of proteins. It is possible to appreciate the different molecular weight of the chimeric protein (pmCherry-hERG-WT) due to the fusion with the fluorescent protein mCherry. β-Actin was used as loading control. (B) Analysis of the cellular distribution of hERG proteins in HEK293 cells. In particular, hERG-mcherry is localized on the plasma membrane as well as the unmodified hERG (pCMV6-XL4-hERG-WT). The mCherry empty vector was used as negative control, the mCherry is distributed homogeneously in all cell compartments and nucleoli.

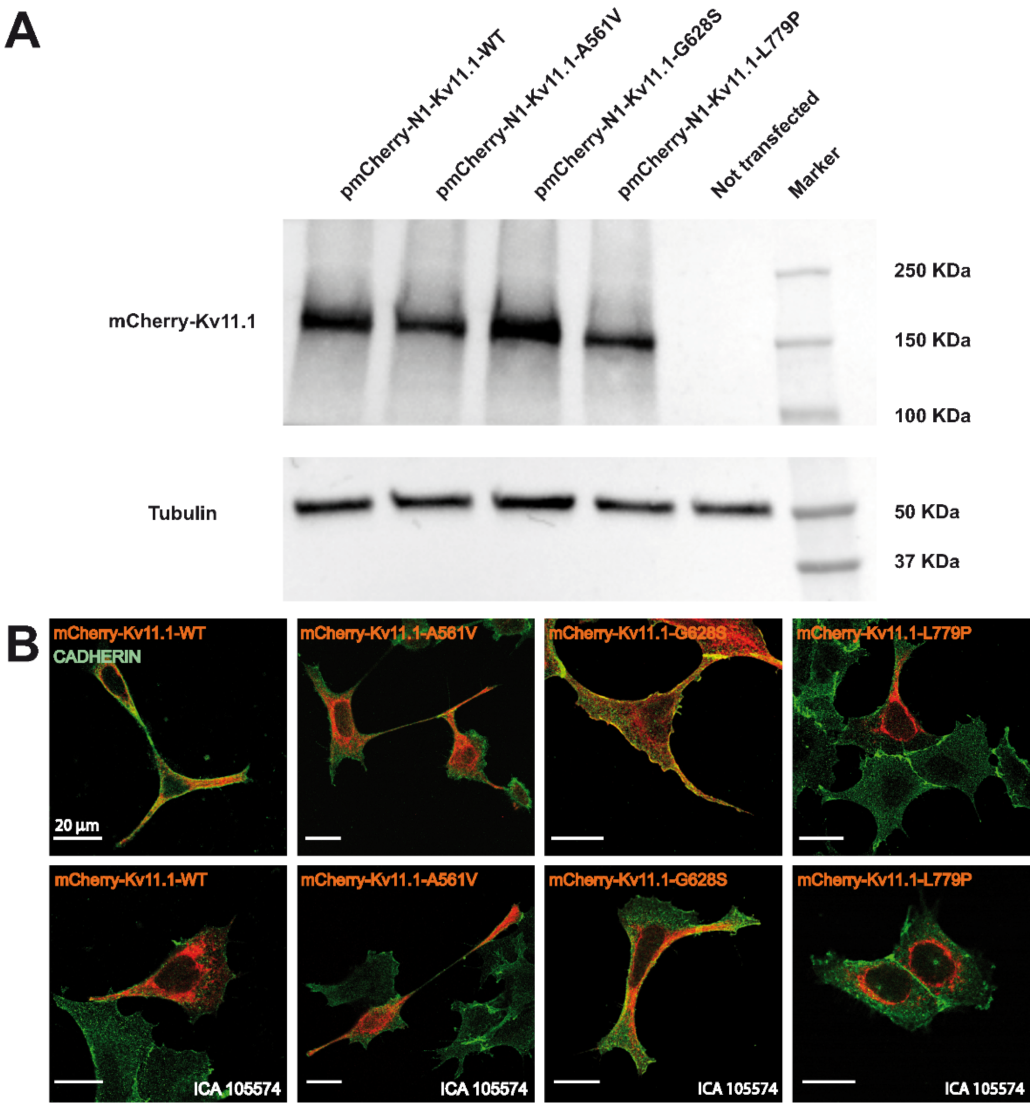

**Figure S4.** Analysis of protein expression of hERG-WT and mutants (-A561V; -G628S and -L779P) after transient transfection in HEK293 cells. (A) Western blot demonstrates the expression of all the proteins. Not transfected cells as negative control and Tubulin was used as loading control. (B) Analysis of cellular localization of pmCherry hERGs (-WT and mutants, respectively) after transfection in HEK293 cells (red) and counterstaining with Ab anti-Cadherin (green) to visualize the plasma membranes. The physiological distribution of hERGs (upper panel) *vs* hERGs localization after ICA-105574 treatment for 2h at 37°C (lower panel). ICA-105574 treatment seems not to affect the hERG distribution, both for the WT and the mutants. All images are visualized by a 63X objective, scale bar 20 µm.

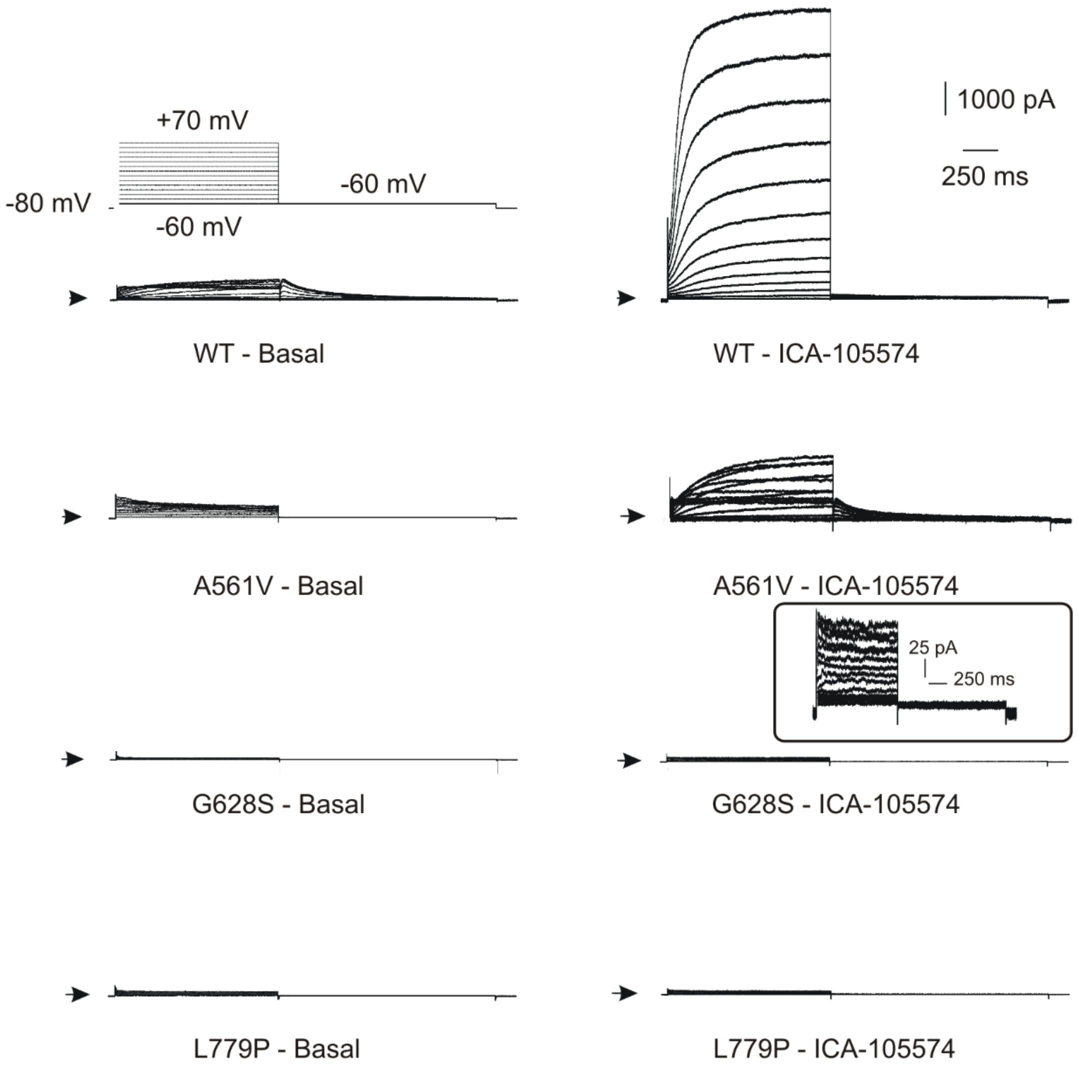

**Figure S5.** Electrophysiological recordings in HEK293 stable cell lines expressing hERG-WT and mutations (-A561V, -G628S, -L779P). The hERG currents were elicited in HEK293 stable cell line by the protocol indicated on the left part of the figure. hERG currents were activated by the depolarizing steps between -60 and +70 mV with a delta of 10 mV, from a holding potential of -80 mV. Each cell was then clamped at -60 mV. In each voltage clamp recording the arrows indicates the zero-current level. The left column reports the voltage clamp recording in resting condition (Resting condition= before the perfusion of the $I_{Kr}$ activator). The right column shows the recordings after the perfusion of ICA-105574 (10 μM). The scale bars reported on the right top are the same for all traces, the inset report more specific scale bars for the hERG G828S mutation perfused with ICA-105574.

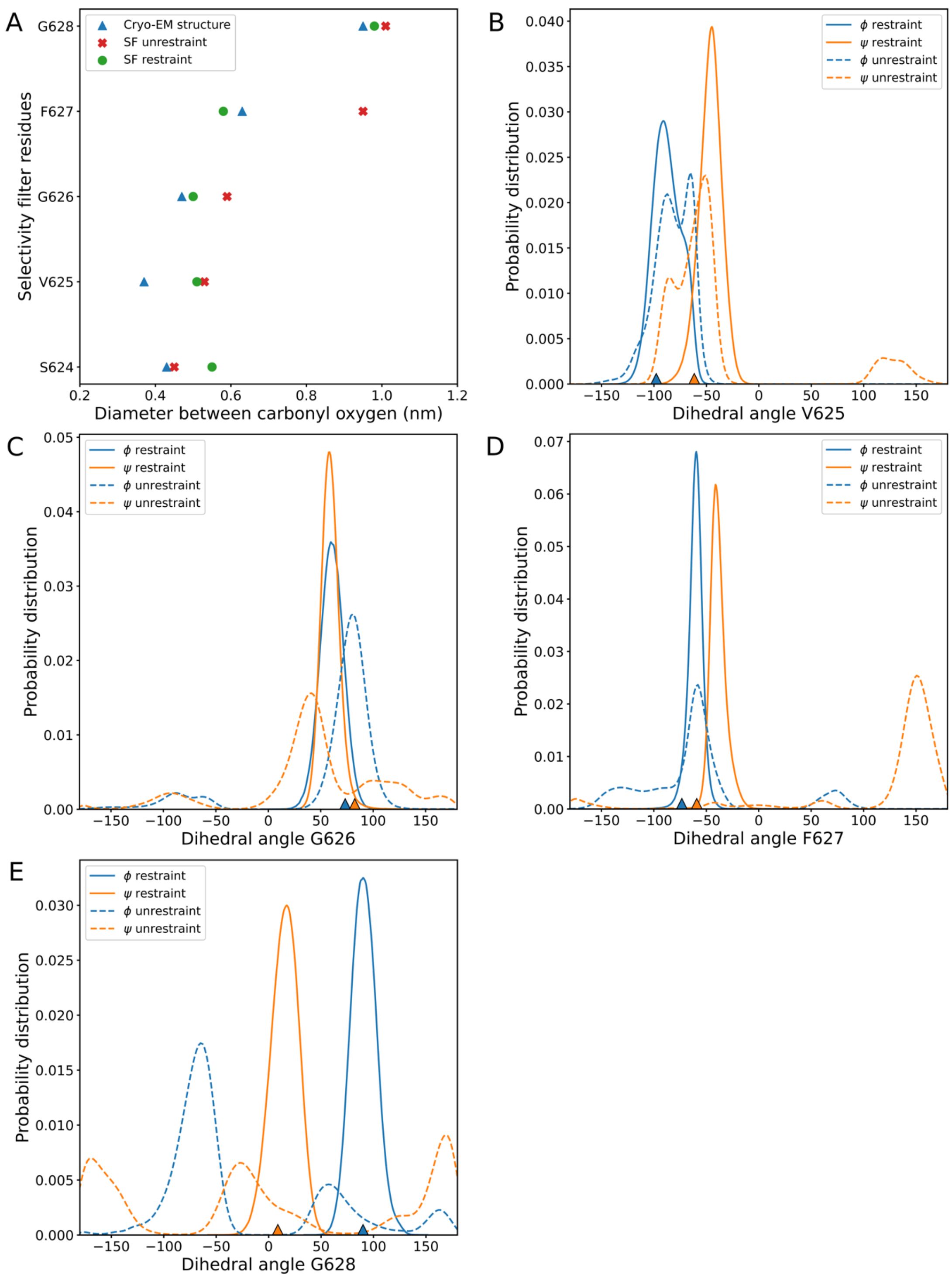

**Figure S6.** The plot demonstrates the necessity to use the dihedral restraints given the fast inactivation and distortion of SF manifested in terms of change in the diameter of the SF pore and backbone dihedrals as highlighted in earlier studies for hERG channels. (A) Comparison of the SF diameter between cryo-EM structure, unrestrained, and restrained MD simulations. In contrast to the unrestrained simulations, which demonstrate an increase in the SF diameter at the extracellular side due to significant fluctuations in the F627 side chains, the restrained simulations show minimal relaxation in the SF while maintaining the overall stability of the pore. (B-E) The probability distribution of dihedral angles (φ, Φ) for the SF residues from the preliminary simulations with and without dihedral restraints. The dihedral values from the cryo-EM structure are marked as triangles for the reference.

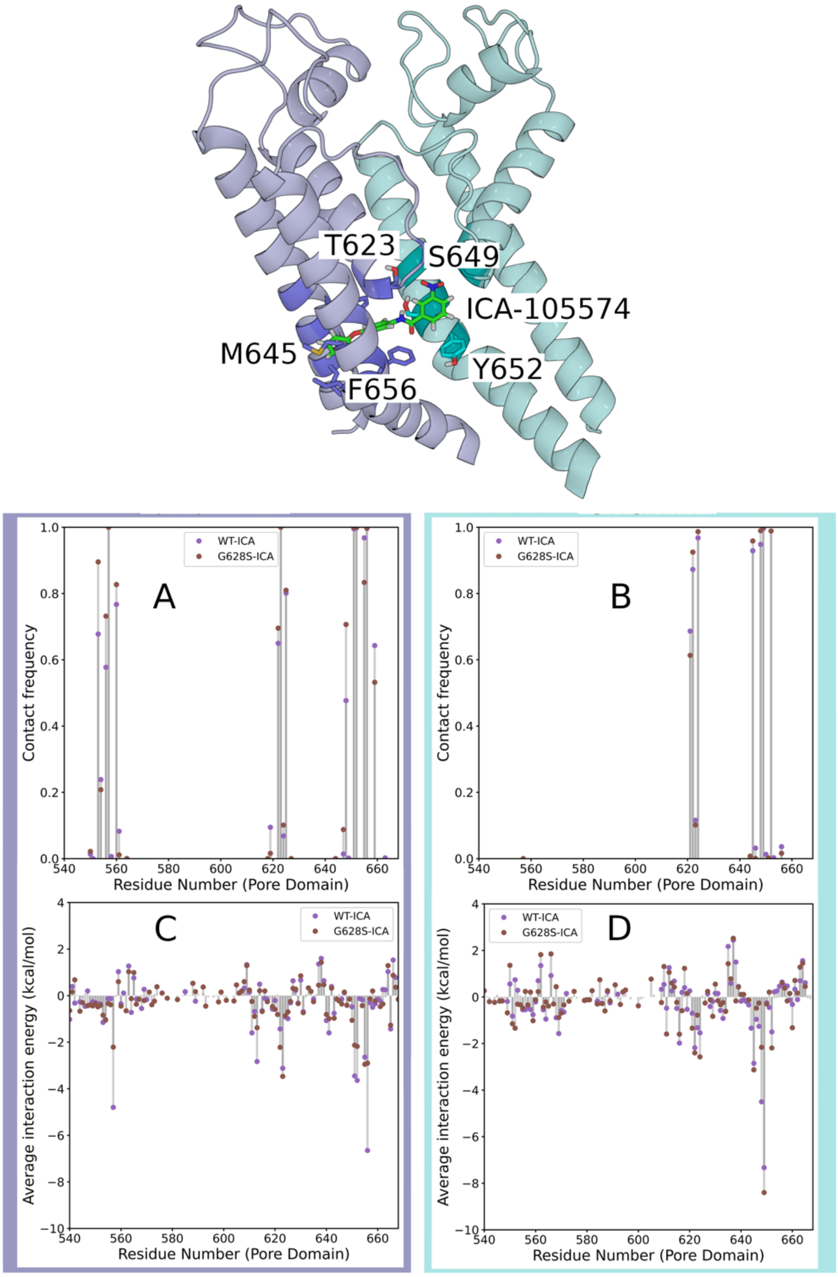

**Figure S7.** Docking pose illustrations of the ICA-105574 molecule between two subunits, with some residues highlighted for position reference. (A, B) Contact map for the interaction of ICA-105574 with different residues in the two opposite subunit fenestrations. (C, D) Average electrostatic interaction energy map (Coulomb + van der Waals) between the ICA-105574 molecule and the pore domain residues. Interestingly, certain residues near the binding site show higher values than other residues.

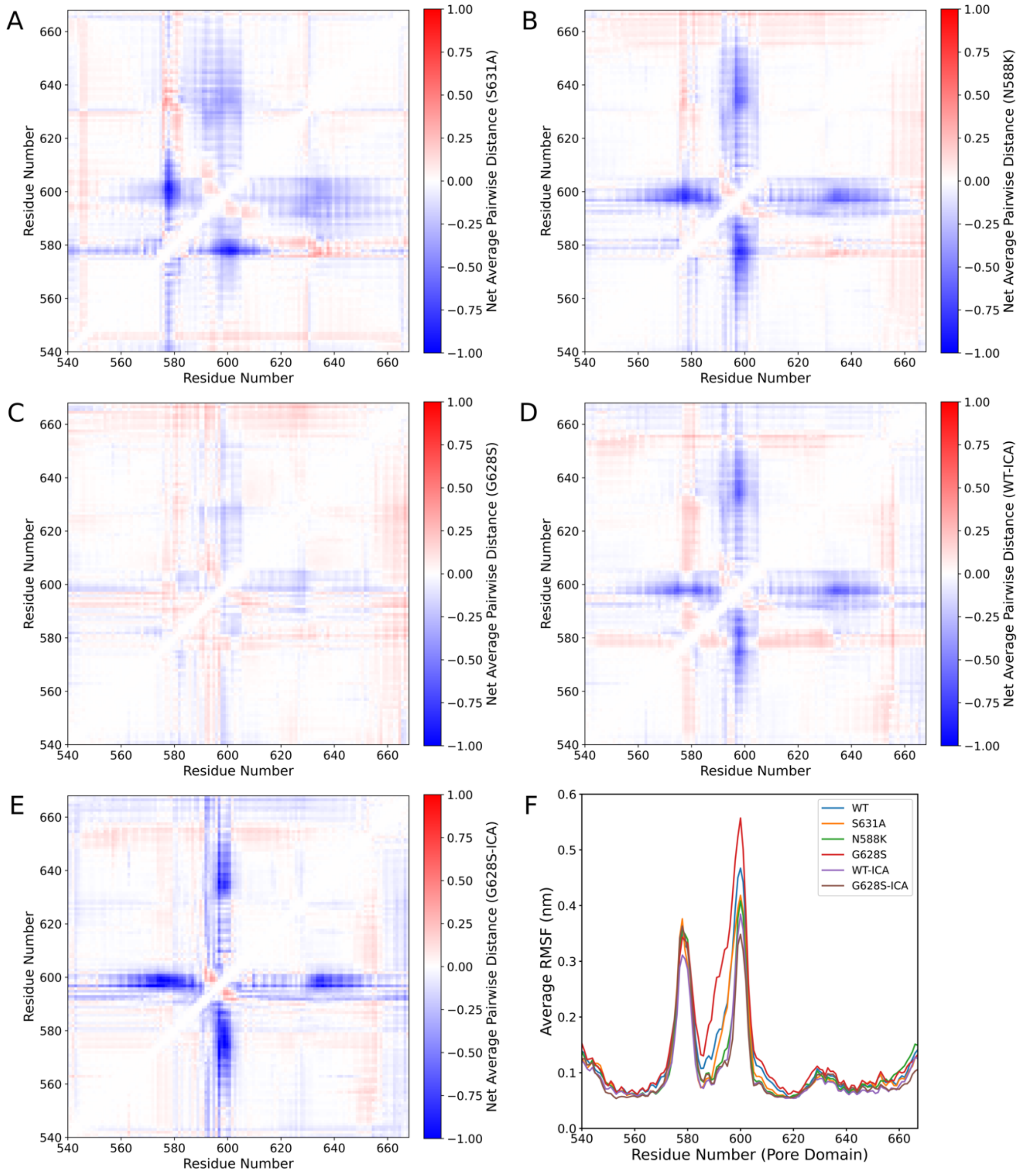

**Figure S8.** (A-E) The figure illustrates the inter-residue pairwise distances for different systems relative to the WT. The net average distance is calculated as the difference in the center of mass (COM) of each residue pair within the pore domain for individual chains between the mutant/ICA-bound systems and WT. Subsequently, we average the pairwise distance for each residue pair across all four chains over all replica trajectories for each system. Our results suggest no significant conformational change in the pore domain. (F) The plot represents root mean square fluctuations (RMSF) calculated for the pore domain and averaged across all the chains over all the replica for each system. It can be observed that the extracellular loops and S5P turret region show higher flexibility in the WT and G628S as compared to the other systems.

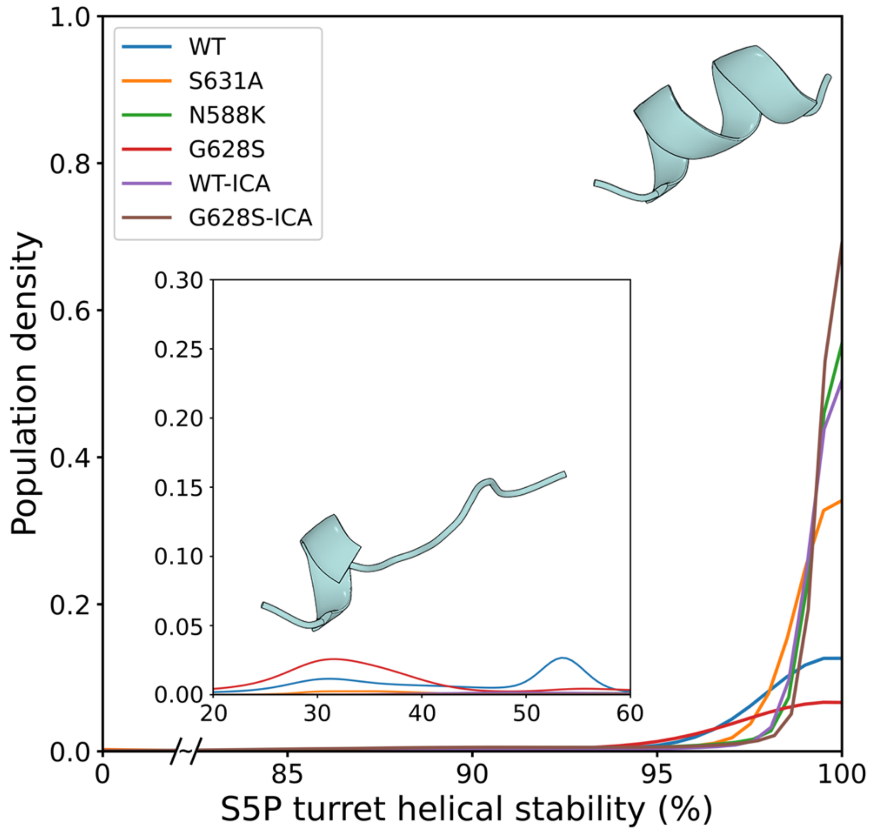

**Figure S9.** Helical propensity of S5P turret region in different systems.

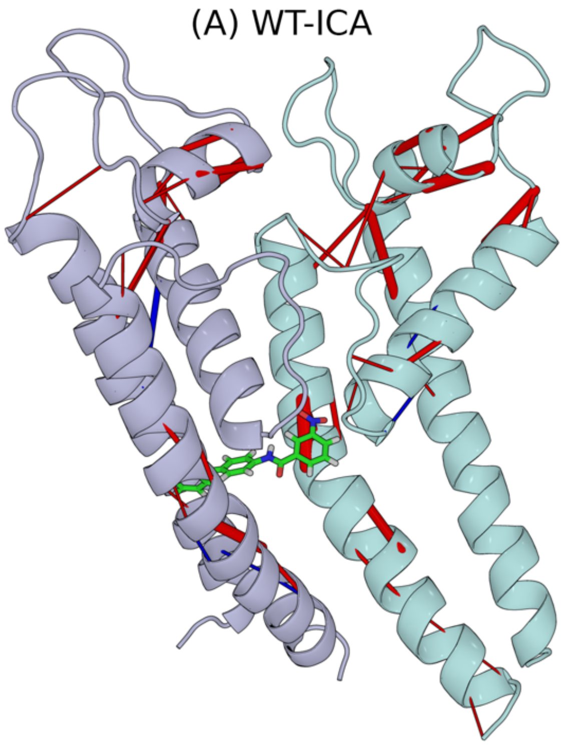

**Figure S10.** Hydrogen bond rearrangement upon perturbation in the form of ICA-105574 binding. The net H-bond occupancy (%) is highlighted in blue and red lines for the reference and the ICA-105574 bound WT system respectively. The thickness of the lines represents the magnitude of the change. (A) Net H-bond occupancy (%) between WT-ICA and WT *(*$\Delta HB_{ij} = HB_{ij}^{WT-ICA} - HB_{ij}^{WT}$*)* shows the effect of ICA-105574. The figure shows two adjacent chains to represent different H-bond patterns following the asymmetric binding of the activator in the hERG. ICA-105574 binding increases H-bond occupancy around SF (G626-S620, S620-Y616) and S5P turret helix (G584-Q592) in both chains, while some residue pairs have opposite H-bond occupancy changes in G628S and ICA-105574 systems (e.g. S641-E637, S649-M645).

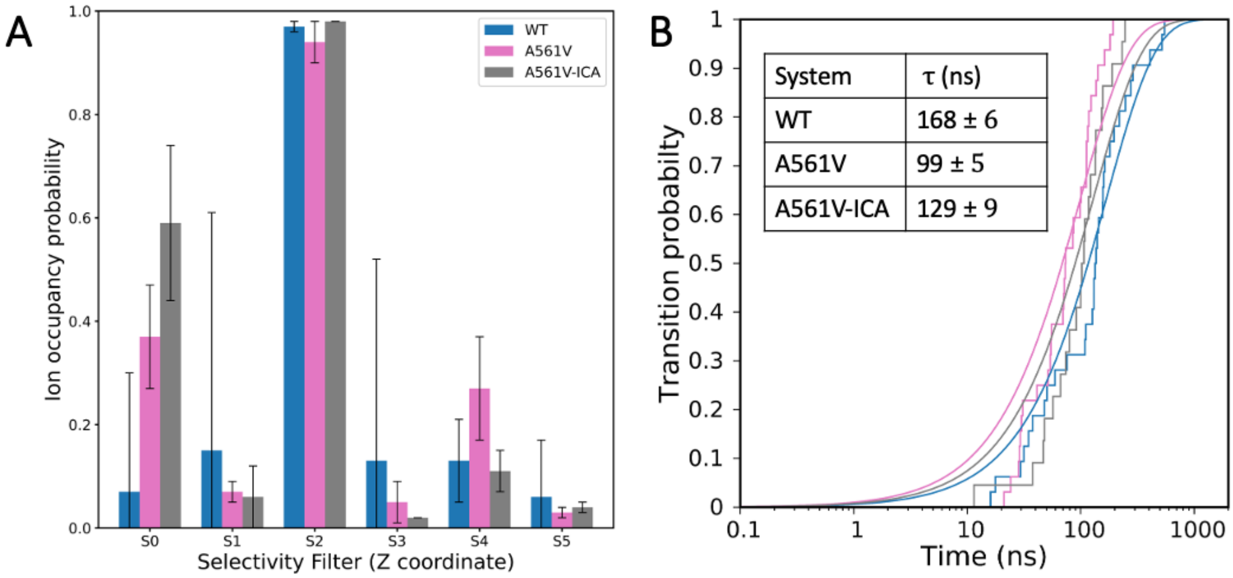

**Figure S11.** (A) The plot represents the ion occupancy probability at different sites in the selectivity filter over the simulation for the A561V and A561V-ICA systems. The selectivity filter is divided into six different sites namely S0 to S4 and S-Cavity. The ion occupancy for both the system is higher at S2 site which in turn prevents the collapse of SF. (B) The plot describes the transition probability of ion crossing event and average transition time for a single permeation event. The inset table shows average transition time for ion permeation event in hERG channel. As observed, the transition time for A561V and A561V-ICA is lower than WT, which could further highlight the limitation of our method in quantifying the conductance and compare directly to the *in vitro* current recordings.

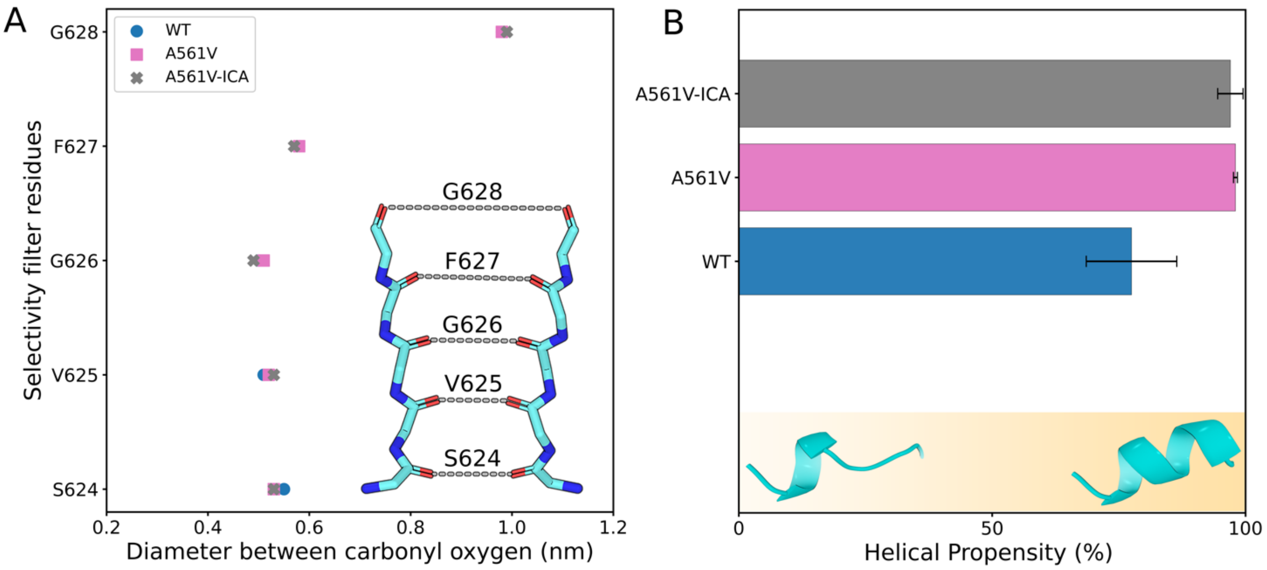

**Figure S12**. (A) The plot shows the diameter of the SF averaged over all replica trajectories for the WT, A561V and A561V-ICA. Unlike G628S mutation, A561V has no adverse effect on the diameter of the SF. (B) The plot shows the helical propensity (%) of S5P turret helix (residue number 584 to 592). The higher helical propensity (> 90%) for A561V and A561V-ICA represents well-structured helix as compared to the WT system. This further highlights the signature of a conducting system.

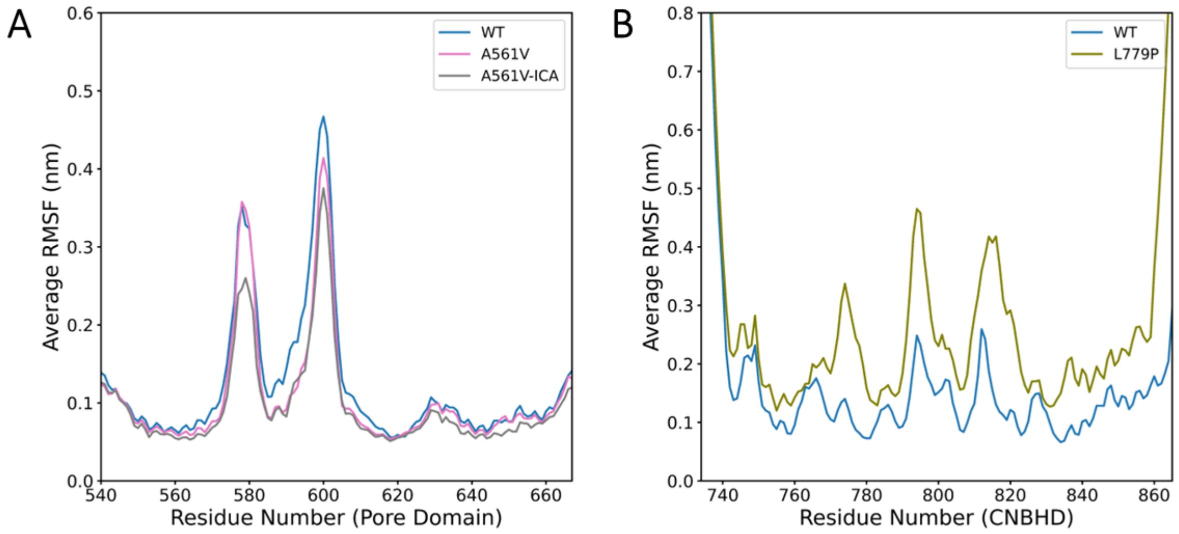

**Figure S13.** (A) The plot represents comparison of RMSF calculated for the pore domain and averaged across all the chains over all the replica between WT, A561V and A561-ICA systems. It can be observed that the extracellular loops and S5P turret region show reduction in flexibility upon the binding of ICA-105574. (B) The average RMSF for the CBNH-domain for simulations performed for wild-type and L779P. The presence of L779P mutation increases the fluctuations in the overall domain suggesting loss of stability of this domain.

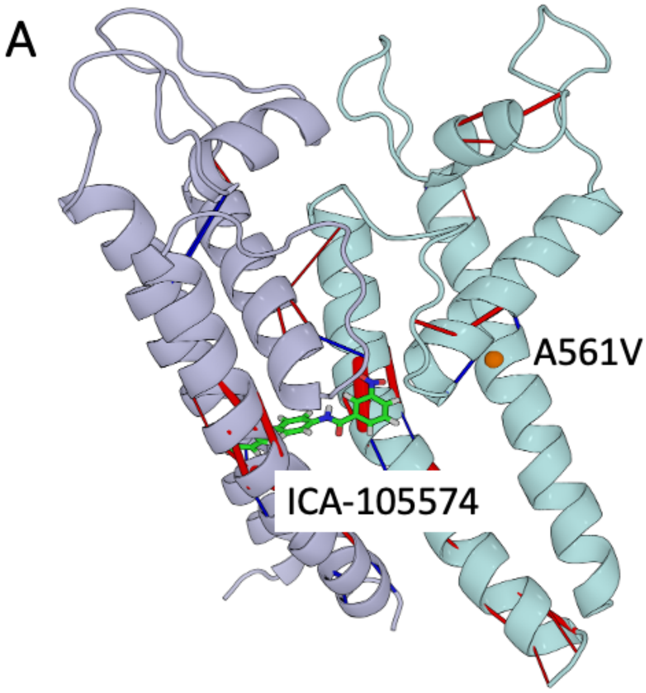

**Figure S14.** Hydrogen bond rearrangement upon A561V mutation or ICA-105574 binding. The net H-bond occupancy (%) is highlighted in blue and red lines for the reference and the perturbed system respectively. The thickness of the lines represents the magnitude of the change. A561V mutation is represented as orange sphere. (A) Net H-bond occupancy (%) between A561V-ICA and A561V ($\Delta HB_{ij} = HB_{ij}^{A561V-ICA} - HB_{ij}^{A561V}$) shows the rescue effect of ICA-105574: The figure shows two adjacent chains to represent different H-bond patterns following the asymmetric binding of the activator in the hERG. ICA-105574 binding induces H-bond rearrangement around SF and S5P turret helix (G584-Q592) in different subunits.

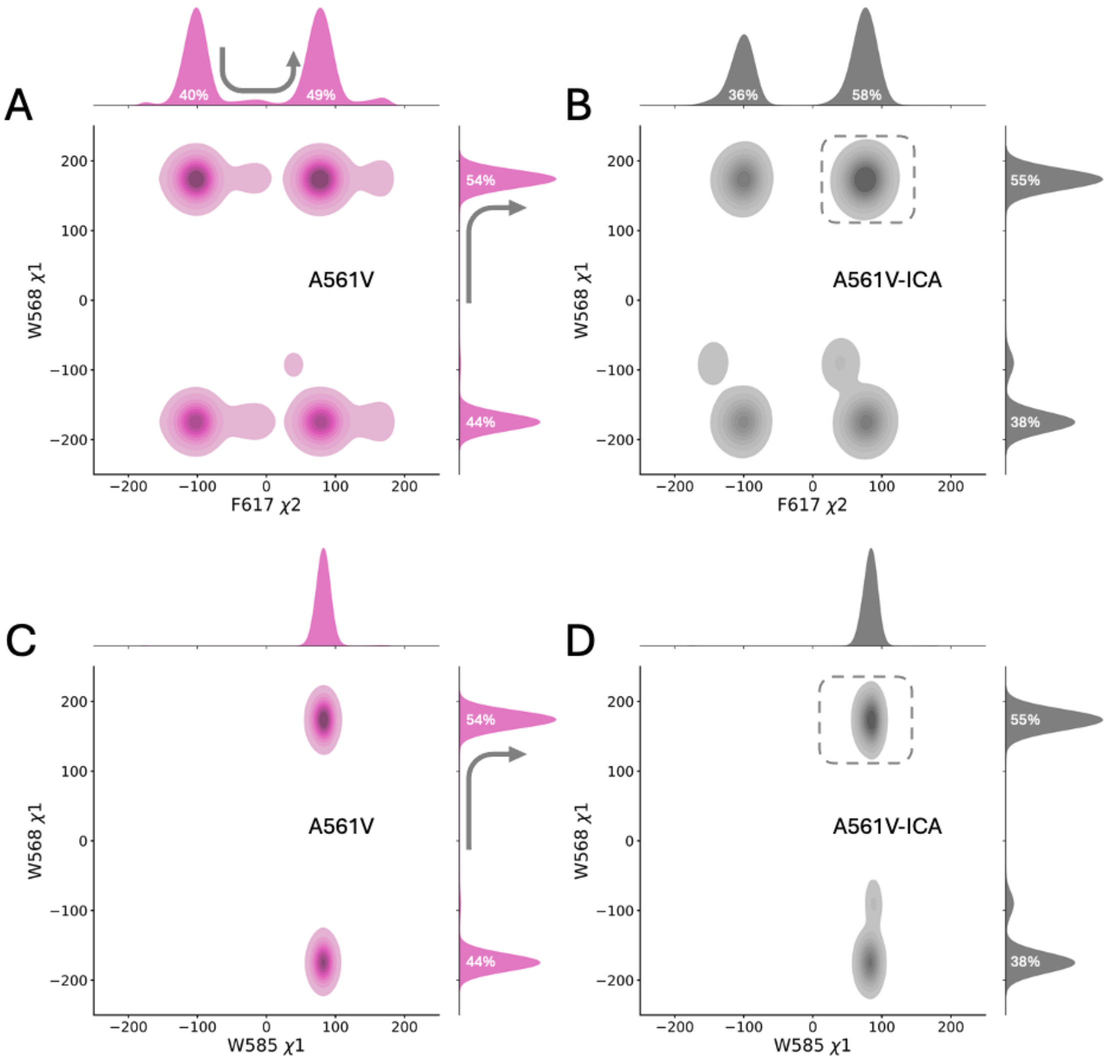

**Figure 15.** (A-D) χ2 dihedral distributions of residue pairs W568-F617 and W568-W585 in the A561V and A561V-ICA systems. The color opacity indicates the population density of each bin. The figure is plotted jointly for each axis variable with a 1D histogram to represent the population shift in the dihedrals. The arrows indicate the changes in the distribution upon ICA-105574 binding, while the dashed square box indicates the most populated basin in 2D distribution upon ICA-105574 binding. (A, B) show the 2D distribution of W568-χ1 and F617-χ2 angles, and (C, D) show the 2D distribution of χ1 angle for W585-W568.

### Supplementary tables

### Table S1. List of preliminary simulations performed to access the stability of selectivity filter.

| *Unrestrained preliminary simulations* | | | | |
|---|---|---|---|---|
| *Systems* | *Replica* | *Production time (ns)* | *Number of ion permeations* | *Membrane Potential (mV)* |
| *WT* | *1* | *1041* | *1* | *~350* |
| | *2* | *812* | *1* | |
| | *3* | *641* | *1* | |
| | *4* | *1035* | *-* | *~700* |
| | *5* | *781* | *1* | |
| | *6* | *599* | *-* | |
| *Dihedral restrained preliminary simulations* | | | | |
| *Systems* | *Ion configuration* | *Replica Number* | *Production time (ns)* | *Membrane Potential (mV)* |
| *WT* | *[KWKWK]* | *1* | *1200* | *~ 700 - 750* |
| | *[KWKWK]* | *2* | *1000* | |
| | *[KKKKK]* | *1* | *900* | |
| | *[KKKKK]* | *2* | *1000* | |
| *Mix Salt (200mM KCl + 200mM NaCl)* | *[KWKWK]* | *1* | *1600* | |
| | *[KWKWK]* | *2* | *1600* | |
| | *[KWKWK]* | *3* | *1600* | |

**Table S2. List of simulations performed and analyzed with dihedral restrain, ion configuration [KWKWK] and 200 mM KCl salt concentration for different systems.**

| *Systems* | *Replica Number* | *Production time (ns)* | *Number of $K^+$ ion permeations* | *Ion transition time (ns)* | *Conductance (pS) at 750mV* |
|---|---|---|---|---|---|
| *WT* | *1* | *2000* | *25* | 168 ± 6 | 1.27 ± 0.04 |
| | *2* | *1000* | *12* | | |
| *S631A* | *1* | *1600* | *24* | 102 ± 3 | 2.10 ± 0.06 |
| | *2* | *1000* | *25* | | |
| *N588K* | *1* | *2000* | *10* | 260 ± 25 | 0.82 ± 0.07 |
| | *2* | *2000* | *12* | | |
| *G628S* | *1* | *1000* | *2* | -- | -- |
| | *2* | *1000* | *1* | | |
| *WT-ICA* | *1* | *1070* | *18* | 128 ± 7 | 1.67 ± 0.09 |
| | *2* | *1000* | *10* | | |
| *G628S-ICA* | *1* | *2000* | *8* | *124 ± 4* | 1.72 ± 0.05 |
| | *2* | *700* | *9* | | |
| *A561V* | *1* | *1000* | *12* | 99 ± 5 | 2.16 ± 0.11 |
| | *2* | *970* | *24* | | |
| *A561V-ICA* | *1* | *1000* | *15* | 129 ± 9 | 1.66 ± 0.12 |
| | *2* | *1000* | *11* | | |

**Table S3. Equilibration protocol details for NVT and NPT ensembles.**

| *Steps* | *Ensemble* | *Time (ns)* | *Position restraint (kJ/mol nm$^2$)*<br>*BB: Backbone, SC: Sidechain*<br>*POPC: Bilayer, DIH: Dihedral restraint on POPC* |
|---|---|---|---|
| *1* | *NVT* | *0.25* | *BB:4000, SC:2000, POPC:1000, DIH:1000, ICA:1000* |
| *2* | *NVT* | *0.5* | *BB:4000, SC:2000, POPC:500, DIH:500, ICA:1000* |
| *3* | *NPT* | *1* | *BB:4000, SC:2000, POPC:400, DIH:200, ICA:1000* |
| *4* | *NPT* | *3* | *BB:4000, SC:2000, POPC:200, DIH:200, ICA:1000* |
| *5* | *NPT* | *3* | *BB:4000, SC:2000, POPC:40, DIH:100, ICA:500* |
| *6* | *NPT* | *5* | *BB:4000, SC:1000, ICA:500* |
| *7* | *NPT* | *5* | *BB:2000, SC:500* |
| *8* | *NPT* | *5* | *BB:1000* |
| *9* | *NPT* | *5* | *BB:500* |

**Table S4. Net H-bond occupancy (%) details from the reference system for perturbations in the form of mutation (G628S) or ligand binding (ICA-105574).** Negative values indicate higher occupancy in the reference system, while positive values indicate higher occupancy in the perturbed system. Net H-bond occupancy (%) for the ligand-bound system is presented for different subunit chains, reflecting the asymmetric binding of the drug. While describing the H-bond between two residues, it is defined as H-bond from Acceptor:Donor with the residue number information and Backbone/Main Chain (M) or Side Chain (S).

| $\Delta HB_{ij} = HB_{ij}^{G628S} - HB_{ij}^{WT}$ | | $\Delta HB_{ij} = HB_{ij}^{G628S-ICA} - HB_{ij}^{G628S}$ | | | |
|---|---|---|---|---|---|
| *N588-M:G584-M* | *-32* | *S654-S:L650-M* | *-31* | *L666-M:I662-M* | *-45* |
| *F656-M:M651-M* | *-27* | *A653-M:S649-M* | *-29* | *Y667-M:I663-M* | *-33* |
| *G626-M:S620-M* | *-26* | *T618-S:A614-M* | *-25* | *R665-M:A661-M* | *-32* |
| *L589-M:W585-M* | *-22* | *Y652-M:G648-M* | *-24* | *I655-M:M651-M* | *-27* |
| *G590-M:L586-M* | *-20* | *I655-M:M651-M* | *-21* | *C566-S:H562-M* | *-25* |
| *N658-M:S654-M* | *-18* | *T623-M:F619-M* | *-20* | *S654-M:L650-M* | *-20* |
| *S621-S:F617-M* | *-16* | *S641-S:E637-M* | *-20* | *T618-S:A614-M* | *-15* |
| *S631-S:N633-S* | *-12* | *S654-M:L650-M* | *-16* | *S641-S:E637-M* | *-15* |
| *S654-S:L650-M* | *-11* | *S649-S:M645-M* | *-14* | *V625-M:S620-M* | *-12* |
| *Y616-S:N629-S* | *13* | *Y569-M:A565-M* | *-12* | *D609-M:S606-S* | *10* |
| *S649-M:M645-M* | *13* | *R541-S:E544-S* | *-10* | *N658-M:S654-M* | *11* |
| *S641-S:E637-M* | *18* | *G594-M:G590-M* | *10* | *W585-M:E637-S* | *12* |
| *I655-M:M651-M* | *18* | *S621-S:F617-M* | *11* | *L666-M:I663-M* | *12* |
| *W585-S:W568-M* | *20* | *L650-M:L646-M* | *11* | *K595-S:D609-S* | *14* |
| *R665-M:A661-M* | *22* | *I662-M:N658-M* | *11* | *S606-S:D609-S* | *15* |
| *Y667-M:I663-M* | *22* | *S606-S:D609-S* | *13* | *S660-S:F656-M* | *16* |
| *L666-M:I662-M* | *27* | *S624-M:S621-M* | *13* | *N598-M:L602-M* | *17* |
| *S606-M:D609-S* | *29* | *L666-M:I662-M* | *14* | *G648-M:V644-M* | *18* |
| *S649-S:M645-M* | *34* | *R665-M:A661-M* | *15* | *S649-M:M645-M* | *20* |
| *K610-M:S606-M* | *40* | *G648-M:V644-M* | *17* | *L650-M:L646-M* | *20* |

| | | | | | |
|---|---|---|---|---|---|
| | | *W585-S:W568-M* | *18* | *S654-S:L650-M* | *20* |
| | | *N598-M:L602-M* | *18* | *S620-S:Y616-M* | *28* |
| | | *S620-S:Y616-M* | *18* | *K595-M:G590-M* | *38* |
| | | *T618-S:A561-M* | *20* | *L589-M:W585-M* | *40* |
| | | *S543-M:D540-M* | *21* | *G590-M:L586-M* | *40* |
| | | *K595-M:G590-M* | *36* | *N588-M:G584-M* | *41* |
| | | *L589-M:W585-M* | *43* | *G626-M:S620-M* | *43* |
| | | *G590-M:L586-M* | *45* | *D591-M:H587-M* | *44* |
| | | *N588-M:G584-M* | *48* | *Q592-M:N588-M* | *57* |
| | | *D591-M:H587-M* | *48* | *I593-M:L589-M* | *66* |
| | | *G626-M:S620-M* | *54* | *F656-M:M651-M* | *76* |
| | | *Q592-M:N588-M* | *60* | | |
| | | *I593-M:L589-M* | *69* | | |

**Table S5. Net H-bond occupancy (%) details from the reference system for perturbations in the form of ligand binding (ICA) or mutations (S631A, N588K) in the SQT1 system.** Negative values indicate higher occupancy in the reference system (WT), while positive values indicate higher occupancy in the perturbed system. Net H-bond occupancy (%) for the ligand-bound system is presented for different subunit chains, reflecting the asymmetric binding of the drug. While describing the H-bond between two residues, it is defined as H-bond from Acceptor: Donor with the residue number information and Backbone/Main Chain (M) or Side Chain (S).

| $\Delta HB_{ij} = HB_{ij}^{WT-ICA} - HB_{ij}^{WT}$ | | | | $\Delta HB_{ij} = HB_{ij}^{S631A} - HB_{ij}^{WT}$ | | $\Delta HB_{ij} = HB_{ij}^{N588K} - HB_{ij}^{WT}$ | |
|---|---|---|---|---|---|---|---|
| *T623-M:F619-M* | *-15* | *Y611-S:H562-S* | *-27* | *L550-M:G546-M* | *-45* | *N588-M:G584-M* | *-58* |
| *T618-S:A614-M* | *-13* | *I663-M:V659-M* | *-16* | *V549-M:Y545-M* | *-39* | *F656-M:M651-M* | *-20* |
| *Q592-S:N588-M* | *10* | *V659-M:I655-M* | *-14* | *F551-M:A547-M* | *-38* | *Q592-M:N588-M* | *-14* |
| *S624-M:S621-M* | *10* | *S654-S:L650-M* | *-13* | *F656-M:M651-M* | *-19* | *G648-M:V644-M* | *10* |
| *L666-M:I662-M* | *10* | *R541-S:E544-S* | *-10* | *S621-S:F617-M* | *-17* | *R665-M:A661-M* | *10* |
| *S631-S:N633-S* | *11* | *S654-M:L650-M* | *-10* | *S631-S:N633-S* | *-17* | *D609-M:S606-S* | *11* |
| *I647-M:C643-M* | *11* | *S606-S:D609-S* | *10* | *L650-M:L646-M* | *-14* | *N573-M:Y569-M* | *12* |
| *I662-M:N658-M* | *11* | *E637-M:T634-S* | *10* | *F627-M:S620-S* | *-10* | *S649-M:M645-M* | *12* |
| *S668-S:R665-M* | *11* | *M651-M:I647-M* | *10* | *Q664-M:S660-M* | *10* | *L666-M:I662-M* | *15* |
| *T618-S:A561-M* | *12* | *L650-M:L646-M* | *11* | *S668-S:R665-M* | *10* | *R582-S:E575-S* | *16* |
| *D609-M:S606-S* | *12* | *R582-S:E575-S* | *12* | *G590-M:L586-M* | *11* | *G590-M:L586-M* | *16* |
| *G626-M:S620-M* | *13* | *N588-M:G584-M* | *13* | *R582-S:E575-S* | *12* | *K595-M:G590-M* | *20* |
| *L589-M:W585-M* | *14* | *G590-M:L586-M* | *13* | *S668-M:R665-M* | *12* | *S606-S:D609-S* | *20* |
| *S631-M:N588-S* | *14* | *W585-M:E637-S* | *17* | *N588-M:G584-M* | *13* | *W585-S:W568-M* | *25* |
| *S668-M:R665-M* | *16* | *S649-M:M645-M* | *19* | *L589-M:W585-M* | *13* | *D591-M:H587-M* | *27* |
| *N658-M:S654-M* | *18* | *S660-S:F656-M* | *20* | *S606-S:D609-S* | *13* | *K588-S:D591-S* | *34* |
| *S606-S:D609-S* | *19* | *G648-M:V644-M* | *22* | *Q592-M:N588-M* | *15* | *S606-M:D609-S* | *38* |
| *W585-M:E637-S* | *21* | *W585-S:W568-M* | *25* | *Q592-S:N588-M* | *16* | *K610-M:S606-M* | *46* |
| *G590-M:L586-M* | *23* | *D591-M:H587-M* | *27* | *D591-M:H587-M* | *19* | *I593-M:L589-M* | *54* |
| *S620-S:Y616-M* | *25* | *S606-M:D609-S* | *32* | *A631-M:N588-S* | *21* | *Q592-M:K588-M* | *59* |

| | | | | | | | |
|---|---|---|---|---|---|---|---|
| *G648-M:V644-M* | *25* | *Q592-M:N588-M* | *37* | *R665-M:A661-M* | *25* | *K588-M:G584-M* | *72* |
| *K595-M:G590-M* | *33* | *K610-M:S606-M* | *43* | *L666-M:I662-M* | *25* | | |
| *S606-M:D609-S* | *38* | *F656-M:M651-M* | *48* | *R541-S:E544-S* | *37* | | |
| *D591-M:H587-M* | *39* | *I593-M:L589-M* | *49* | *S606-M:D609-S* | *38* | | |
| *W585-S:W568-M* | *42* | | | *K610-M:S606-M* | *44* | | |
| *F656-M:M651-M* | *42* | | | *W585-S:W568-M* | *54* | | |
| *K610-M:S606-M* | *44* | | | *W585-M:E637-S* | *55* | | |
| *Q592-M:N588-M* | *48* | | | | | | |
| *I593-M:L589-M* | *57* | | | | | | |
| *S649-S:M645-M* | *64* | | | | | | |

**Table S6. Net H-bond occupancy (%) details from the reference system for perturbations in the form of mutation (A561V) or ligand binding (ICA-105574).** Negative values indicate higher occupancy in the reference system, while positive values indicate higher occupancy in the perturbed system. Net H-bond occupancy (%) for the ligand-bound system is presented for different subunit chains, reflecting the asymmetric binding of the drug. While describing the H-bond between two residues, it is defined as H-bond from Acceptor: Donor with the residue number information and Backbone/Main Chain (M) or Side Chain (S).

| $\Delta HB_{ij} = HB_{ij}^{A561V} - HB_{ij}^{WT}$ | | $\Delta HB_{ij} = HB_{ij}^{A561V-ICA} - HB_{ij}^{A561V}$ | | | |
|---|---|---|---|---|---|
| *F656-M:M651-M* | -17 | *M574-M:I571-M* | -24 | *L666-M:I662-M* | -21 |
| *S649-M:M645-M* | -13 | *C566-S:H562-M* | -16 | *W585-S:W568-M* | -18 |
| *L650-M:L646-M* | -11 | *T623-M:F619-M* | -15 | *Y667-M:I663-M* | -16 |
| *Y569-S:E435-S* | -10 | *A653-M:S649-M* | -13 | *V625-M:S620-M* | -14 |
| *G648-M:V644-M* | -10 | *S543-S:L539-M* | -12 | *R665-M:A661-M* | -14 |
| *N573-S:Y569-M* | 11 | *Y542-S:D411-M* | -10 | *S654-M:L650-M* | -13 |
| *D609-M:S606-S* | 11 | *Y652-M:G648-M* | -10 | *V659-M:I655-M* | -13 |
| *V625-M:S620-M* | 12 | *N573-M:Y569-M* | 10 | *Y542-S:D411-M* | -12 |
| *L589-M:W585-M* | 13 | *I647-M:C643-M* | 10 | *W585-M:E637-S* | -12 |
| *S620-S:Y616-M* | 13 | *Y667-M:I663-M* | 11 | *F557-M:L553-M* | -10 |
| *N588-M:G584-M* | 14 | *S668-S:R665-M* | 11 | *Y616-S:N629-S* | 10 |
| *G590-M:L586-M* | 15 | *Q592-M:N588-M* | 14 | *I647-M:C643-M* | 10 |
| *K595-M:G590-M* | 16 | *N658-M:S654-M* | 15 | *S606-S:D609-S* | 11 |
| *S606-S:D609-S* | 19 | *S668-M:R665-M* | 15 | *S621-S:F617-M* | 13 |
| *D591-M:H587-M* | 24 | *G626-M:S620-M* | 17 | *S606-M:D609-S* | 14 |
| *S606-M:D609-S* | 29 | *S649-M:M645-M* | 17 | *S620-S:Y616-M* | 14 |
| *Q592-M:N588-M* | 33 | *R665-M:A661-M* | 17 | *M651-M:I647-M* | 16 |
| *K610-M:S606-M* | 41 | *L650-M:L646-M* | 18 | *S660-S:F656-M* | 18 |
| *W585-S:W568-M* | 43 | *D591-M:H587-M* | 19 | *L650-M:L646-M* | 30 |
| *I593-M:L589-M* | 47 | *K595-M:G590-M* | 20 | *S649-M:M645-M* | 33 |

| | | | | | |
|---|---|---|---|---|---|
| | | *S620-S:Y616-M* | *22* | *G648-M:V644-M* | *34* |
| | | *L666-M:I662-M* | *27* | *F656-M:M651-M* | *68* |
| | | *G648-M:V644-M* | *35* | | |
| | | *F656-M:M651-M* | *56* | | |
| | | *S649-S:M645-M* | *66* | | |

**Legends for Datasets**

The dataset is a collection of molecular dynamics (MD) simulation trajectories generated using GROMACS software, initial structure (gro and pdb format) and topology files (tpr) used for performing MD simulations to investigate the therapeutic potential of ICA-105574, a hERG activator, in severe hERG mutations (A561V, G628S, L779P) associated with LQT2. Details about different ZIP files:

1. WT: WT.

2. SQT mutations: S631A, N588K.

3. LQT mutations: G628S, A561V.

4. ICA-105574 bound trajectories: WT-ICA, G628S-ICA and A561V-ICA.

5. LQT mutation: L779P (MD simulations using only CNBHD in the WT (WT) and mutation (MUT).

6. MDP-PLUMED: MDP files for running different steps in simulations using GROMACS.

Files in each folder:

1. SYSTEM_Charmm-Gui_Output.pdb : Output file generated using charmm-gui protocol. The file contains: Protein embedded in POPC bilayer with water and KCl (200mM).

2. Structure.gro: Structure after minimisation with $K^+$ placed manually (KWKWK configuration) inside the selectivity filter to prevent inactivation.

3. R1 and R2: Two replica of trajectories. The trajectories are saved with frames at every 1ns for convenience. Full trajectories with frames saved at every 10ps are available upon request.

4 md.tpr: The file represents the topology parameters used for the production run. The number of steps and the starting structure may differ depending upon the individual equilibration and final length of the production run for different replica.

**Legends for Movie S1**

The movie shows a few $K^+$ permeation event from one of the WT-hERG MD simulations. Only two chains of the protein are partially visible and represented as cartoon and surface envelope. $K^+$ ions are shown as colored spheres while water molecules surrounding them are represented in ball-and-stick.

### References

1. Li Y, Ng HQ, Li Q, Kang CB. Structure of the Cyclic Nucleotide-Binding Homology Domain of the hERG Channel and Its Insight into Type 2 Long QT Syndrome. Scientific Reports 2016 6:1 [Internet]. 2016 [cited 2024 Mar 7];6:1–10. Available from: https://www.nature.com/articles/srep23712
